# Genomic changes underlying host specialization in the bee gut symbiont *Lactobacillus* Firm5

**DOI:** 10.1101/483685

**Authors:** KM Ellegaard, S Brochet, G Bonilla-Rosso, O Emery, N Glover, N Hadadi, KS Jaron, JR van der Meer, M Robinson-Rechavi, V Sentchilo, F Tagini, SAGE class 2016-17, P Engel

## Abstract

Bacteria that engage in longstanding associations with particular hosts are expected to evolve host-specific adaptations that limit their capacity to thrive in other environments. Consistent with this, many gut symbionts seem to have a limited host range, based on community profiling and phylogenomics. However, few studies have experimentally investigated host specialization of gut symbionts and underlying mechanisms have largely remained elusive. Here, we studied host specialization of a dominant gut symbiont of social bees, *Lactobacillus* Firm5. We show that Firm5 strains isolated from honey bees and bumble bees separate into deep-branching host-specific phylogenetic lineages. Despite their divergent evolution, colonization experiments show that bumble bee strains are capable of colonizing the honey bee gut. However, they were less successful than honey bee strains, and competition with honey bee strains completely abolished their colonization. In contrast honey bee strains of divergent phylogenetic lineages were able to coexist within individual bees. This suggests that both host selection and interbacterial competition play important roles for host specialization. Using comparative genomics of 27 Firm5 isolates, we found that the genomes of honey bee strains harbor more carbohydrate-related functions than bumble bee strains, possibly providing a competitive advantage in the honey bee gut. Remarkably, most of the genes encoding carbohydrate-related functions were not conserved among the honey bee strains, which suggests that honey bees can support a metabolically more diverse community of Firm5 strains than bumble bees. These findings advance our understanding of genomic changes underlying host specialization.

## Introduction

Symbiotic relationships between bacteria and eukaryotes are pervasive and range from loose associations to obligate interdependencies (McFall-Ngai *et al.* 2013; Kostic *et al.* 2013). The evolution of a host-associated lifestyle is typically accompanied by the loss of generalist characteristics, limiting a symbionts’ capacity to compete and survive in other environments. This in turn results in host specialization (Bobay & Ochman 2017; Sriswasdi *et al.* 2017). In particular, bacteria with longstanding associations are often host-specific and undergo marked genomic changes (Toft & Andersson 2010). Among the most extreme examples are primary endosymbionts of plant-sap feeding insects. These obligate mutualists reside within host cells, are vertically inherited through the germ-line, and have experienced extreme genome reduction due to population bottlenecks and genetic drift (McCutcheon & Moran 2012).

Based on phylogenetic analyses, host specialization has also been inferred for many gut symbionts, as bacterial lineages are frequently found to be exclusively associated with particular hosts (Ley *et al.* 2008; Oh *et al.* 2010; Ochman *et al.* 2010; Eren *et al.* 2015; Moeller *et al.* 2016; Kwong *et al.* 2017). This is remarkable considering that gut symbionts can be horizontally transmitted and are exposed at least at some point to the environment outside the host, which in principle provides opportunities for host switching.

This leads to the question whether the observed host association is determined by differences in the symbiont’s fundamental niches (the set of environmental and nutritional requirements that allow growth in specific hosts), or by constrains due to biological interactions (e.g. cross-feeding, competition, dispersion), i.e their realized (observed) niche is a subset of their fundamental niche (Macarthur & Levins 2015; Hutchinson 1957).

In the case of the gut symbiont *Lactobacillus reuteri,* strains isolated from mice are capable of colonizing the mouse gut, whereas those from humans, pigs, or chickens are not, suggesting that host association in this case has resulted in the restriction of the fundamental niche (Oh *et al.* 2010; Frese *et al.* 2011). In contrast, in another study it was shown that bacterial communities from diverse habitats can colonize and persist in the mouse gut (despite the fact that these species naturally do not occur in the mouse gut), suggesting that a species’ realized niche is frequently more restricted than its fundamental niche (Seedorf *et al.* 2014). A notable difference between the two studies is that host specificity of *L. reuteri* was tested in mice that were free of *Lactobacilli*, but otherwise harbored a conventionalized microbiota, whereas in the second study, most experiments were carried out in microbiota-free mice, supporting the notion that competition can limit the realized niche of gut symbionts (Seedorf *et al.* 2014).

Given that both experimental and phylogenetic evidence is required to determine host specialization, our understanding of host specialization is limited for most gut symbionts. Moreover, little is known about the underlying mechanisms and the genomic changes accompanying host-specific evolution of gut symbionts. Selective forces acting on gut symbionts may differ between hosts due to varying degrees of population bottlenecks during transmission, or due to differences in dietary preferences, gut structure or host physiology, resulting in distinct evolutionary patterns.

A good model to study host specialization of bacterial inhabitants is the gut microbiota of corbiculate bees (Kwong & Moran 2015). Most species of honey bees, bumble bees, and stingless bees share a specialized core gut microbiota that is composed of five phylotypes (strains sharing ≥97% 16S rRNA sequence identity as estimated from amplicon sequencing studies): the gammaproteobacterium *Gilliamella*, the betaproteobacterium *Snodgrassella alvi,* two Lactobacilliales (Firm5 and Firm4), and a Bifidobacterium (Cox-Foster *et al.* 2007; Moran *et al.* 2012; Corby-Harris *et al.* 2014). These phylotypes are likely to have been acquired in a last common ancestor of the corbiculate bees, as they are widely distributed among contemporary species of honey bees, bumble bees, and stingless bees (Kwong *et al.* 2017). Moreover, there is evidence for host specialization and coevolution, because strains isolated from the three groups of corbiculate bees separate into divergent sublineages for most phylotypes (Koch *et al.* 2013; Kwong *et al.* 2014; Ellegaard *et al.* 2015; Zheng *et al.* 2016; Kwong *et al.* 2017; Steele *et al.* 2017).

The best-studied member of the bee gut microbiota with respect to host specialization is *S. alvi* (Kwong & Moran 2015). Reciprocal mono-colonization experiments of microbiota-depleted bees showed that *S. alvi* isolates from honey bees *(Apis mellifera)* colonize the gut of bumble bees *(Bombus impatiens)* poorly, and vice versa, suggesting that the host-specific evolution of these isolates has led to specialization (Kwong *et al.* 2014). Based on the comparison of three *S. alvi* genomes, it was suggested that bumble bee isolates tend to have smaller genomes and contain larger amounts of mobile elements than honey bee isolates. Genomic differences were also identified among isolates from different host groups (honey bees and bumble bees) for the phylotype *Gilliamella:* honey bee isolates encoded more carbohydrate-related functions than bumble bee isolates (Kwong *et al.* 2014). However, recent genome sequencing of a larger number of *Gilliamella* strains revealed that some isolates from honey bees have genomes as small as those from bumble bees, suggesting that a large metabolic repertoire is not strictly needed for colonization of the honey bee gut (Zheng *et al.* 2016; Ludvigsen *et al.* 2017; Steele *et al.* 2017).

For the other phylotypes of the bee gut microbiota, little is known about the link between phylogeny, host range, and genome features. One of the most widely distributed and abundant phylotypes of the bee gut microbiota is *Lactobacillus* Firm5. In the gut of honey bees *(Apis mellifera),* four deep-branching sublineages have been identified for this phylotype (Ellegaard et al. 2015), with pairwise average nucleotide identity (ANI) values well below 90% across sublineages (Ellegaard & Engel 2019), despite their relatively conserved 16S rRNA sequences (≥ 96.5% identity).

Overall, strains of different sublineages were found to vary up to 40% in gene content (Ellegaard et al. 2015), suggesting that they may have adapted to distinct metabolic or spatial niches within the honey bee gut, and leading to the proposition of different species names (Olofsson *et al.* 2014). Interestingly, Firm5 strains isolated from other corbiculate bees seem to belong to different sublineages than the honey bee isolates, as indicated by single gene phylogenies (Kwong *et al.* 2017). Moreover, a divergent Firm5 strain from bumble bees has been isolated and described as a new species, *Lactobacillus bombicola* (Praet *et al.* 2015). Given that Firm5 strains can be cultured, and that the honey bee is amenable to experimental colonization, this phylotype represents an excellent opportunity to study evolutionary trajectories of host adaptation and the consequences for the fundamental and realized niche of this gut symbiont.

Here, we used experimental colonization, genome sequencing, and comparative genomics to address host specialization in the Firm5 phylotype. First, we show that isolates from honey bees and bumble bees belong to distinct sublineages, suggesting longstanding host-specific associations. Second, we provide experimental evidence that both host selection and interbacterial competition contribute to host specialization. Third, our comparative genome analysis reveals marked differences in carbohydrate utilization capacities between honey bee and bumble bee isolates, suggesting that adaptation to the honey bee gut has resulted in larger metabolic flexibility than adaptation to the bumble bee gut.

## Results

### Bumble bee and honey bee isolates belong to separate sublineages of the Firm5 phylotype

We sequenced the genomes of 15 new isolates of the Firm5 phylotype. Five isolates were obtained from the honey bee *Apis mellifera* and ten isolates from three different bumble bee species (five from *Bombus pascuorum,* four from *Bombus bohemicus,* and one from *Bombus terrestris).* All bees were collected in Western Switzerland (Table S1). We also included 12 previously sequenced isolates (one from a bumble bee, the others from honey bees) to be more comprehensive in our analyses. All 27 isolates shared >95% sequence identity across the full-length 16S rRNA gene (**Table S2**), with isolates from conspecific hosts having higher 16S rRNA sequence identities in most cases. The draft genomes of the 15 newly sequenced isolates consisted of 11-24 contigs with total lengths of 1.63-2.11 Mb, which is in the range of the previously sequenced Firm5 strains (**Table S1**). While the genomes of the bumble bee isolates tended to be smaller (1.63-1.70 Mb) than those of the honey bee isolates (1.68-2.15 Mb) (Kruskal-Wallis: chi=14.8, d.f.=1, p-value = 0.0001186), genome synteny was largely conserved across the entire Firm5 phylotype (**Figure S1 and S2**).

To assess the evolutionary relationship between the 27 sequenced Firm5 strains, we inferred a genome-wide phylogeny (including 15 close and three more distant outgroup strains, see methods) (**Figure 1**), and calculated pairwise average nucleotide identities (ANI) (**Figure S3, Table S3**). These analyses showed that the Firm5 strains fall into six monophyletic sublineages with >96% ANI for within-lineage divergence in all cases except “Firm5-4”, for which pairwise ANI values were as low as 91%. All ANI values were <86% between sublineages, indicating the presence of a discontinuity zone between 86 and 91% (Jain et al. 2018), and suggesting that the sublineages correspond to distinct species. Four of these six sublineages consisted of only honey bee isolates and corresponded to the previously identified Firm5 sublineages (Ellegaard *et al.* 2015). The two other sublineages consisted of only bumble bee isolates and formed a monophyletic clade within Firm5 (**Figure 1**). One sublineage comprised isolates from three different bumble bee species *(B. lapidarus, B. terrestris, B. pascuorum)* including the previous isolate described as species *L. bombicola* (Praet *et al.* 2015). The other sublineage comprised exclusively isolates from *B. bohemicus*. Based on its deep divergence from the other sublineages (ANI <80%, **Figure S3**), this second sublineage of bumble bee isolates is likely to represent a novel species.

**Figure 1.**
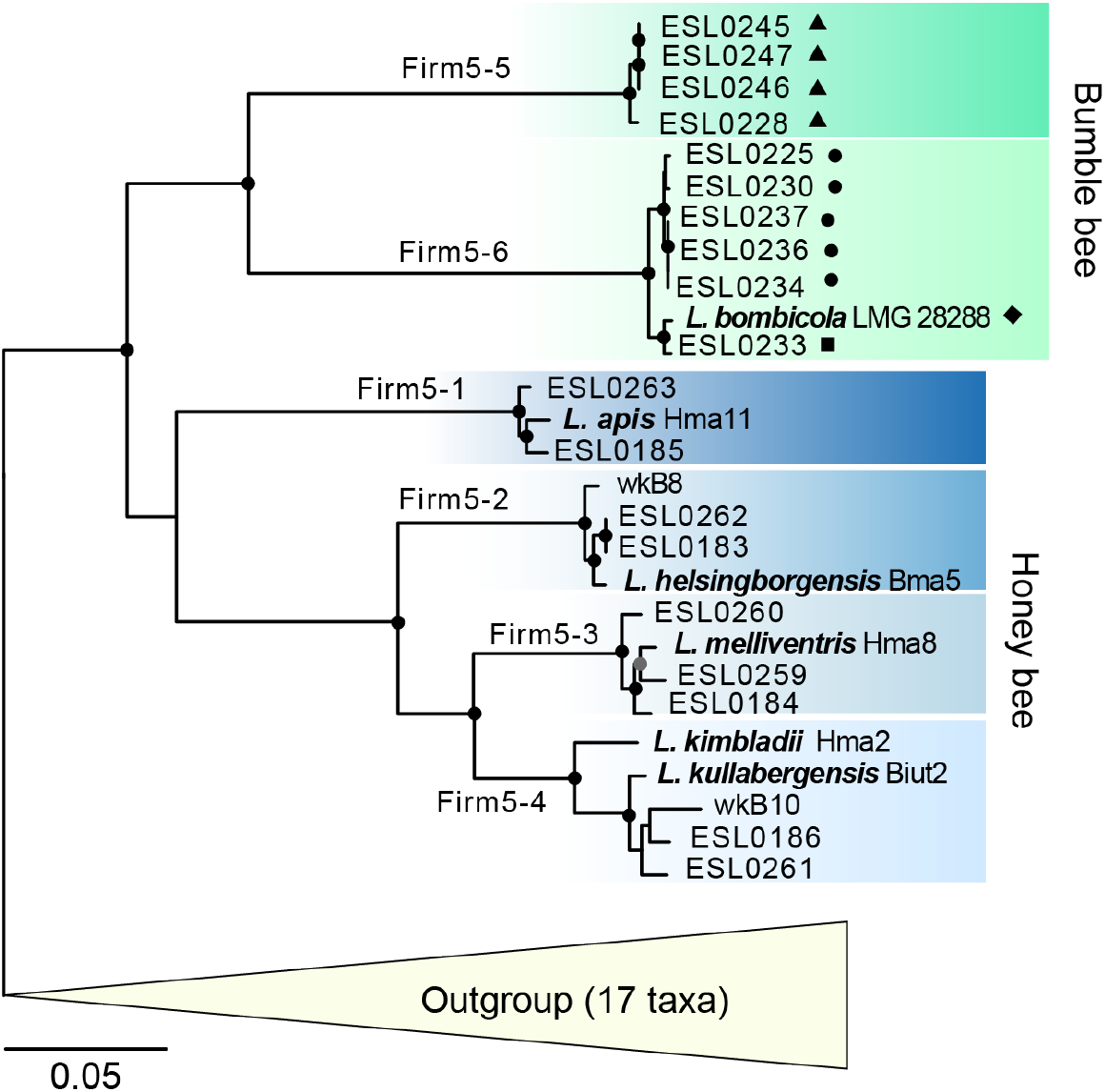
Core genome phylogeny of *Lactobacillus* Firm5. The tree was inferred using maximum likelihood on the concatenated protein alignments of 408 single-copy core genes (i.e. present in all Firm5 strains and in the outgroup strains). The collapsed outgroup consisted of 17 strains that were used to root the tree (see **Figure S4** for the complete tree). The two lineages of bumble bee strains and the four lineages of honey bee strains are shown in green and blue color shades, respectively. Black and grey circles indicate bootstrap support values of 100 and ≥80, respectively, out of 100 replicates. The strain designation of each isolate is given and the species names of the type strains are indicated. The length of the bar indicates 0.05 amino acid substitutions/site. Shapes behind strain name of bumble bee isolate indicate the bee species that the strain was isolated from. Triangle, *Bombus bohemicus;* circle, *Bombus pascuorum;* rhombus, *Bombus lapidarius;* square, *Bombus terrestris.*

Out of the 27 Firm5 isolates included in the current study, five isolates (ESL0262, ESL0234, ESL0236, ESL0245, ESL0247) from three different sublineages were identical or almost identical to other isolates (ANI >99.99%, **Table S3**). In all cases, the nearly identical isolates were obtained from the same individual. Hence, they were excluded from all subsequent analyses to avoid biases due to repeated sampling of the same genotype.

In summary, our phylogenetic analysis of the Firm5 phylotype revealed a pattern suggesting host specialization, because strains of each of the six deep-branching sublineages were exclusively associated with either honey bees or bumble bees.

### Bumble bee strains can colonize microbiota-depleted honey bees, but are outcompeted by honey bee strains

To test whether the fundamental niche of the bumble bee strains extends to the honey bee (*A. mellifera*) we experimentally tested the ability of bumble bee strains to colonize the honey bee gut. In the absence of competitors (i.e. when microbiota-depleted bees were mono-colonized), we found that all bumble bee strains were able to colonize the honey bee hindgut (**Figure 2A**). The number of recovered bacterial cells at day 5 post colonization (10^5^ – >10^8^ CFUs per gut) was substantially higher than in the inoculum (**Figure S5A**) indicating active growth of the bumble bee strains in the honey bee gut. The fact that they can successfully colonize and grow means that the fundamental niche of the bumble bee strains also includes the honey bee gut. However, the percentage of successfully colonized bees was lower for bumble bee strains (10-80%) than for honey bee strains (80-100%) (chi=14.1, d.f.=1, p = 0.0002, and colonization efficiency was slightly lower compared to mono-colonizations with honey bee strains, which reached 10^7^-10^9^ CFUs per gut (**Figure 2A**) (F=38.1, d.f=1, p = 0.0008).

**Figure 2.**
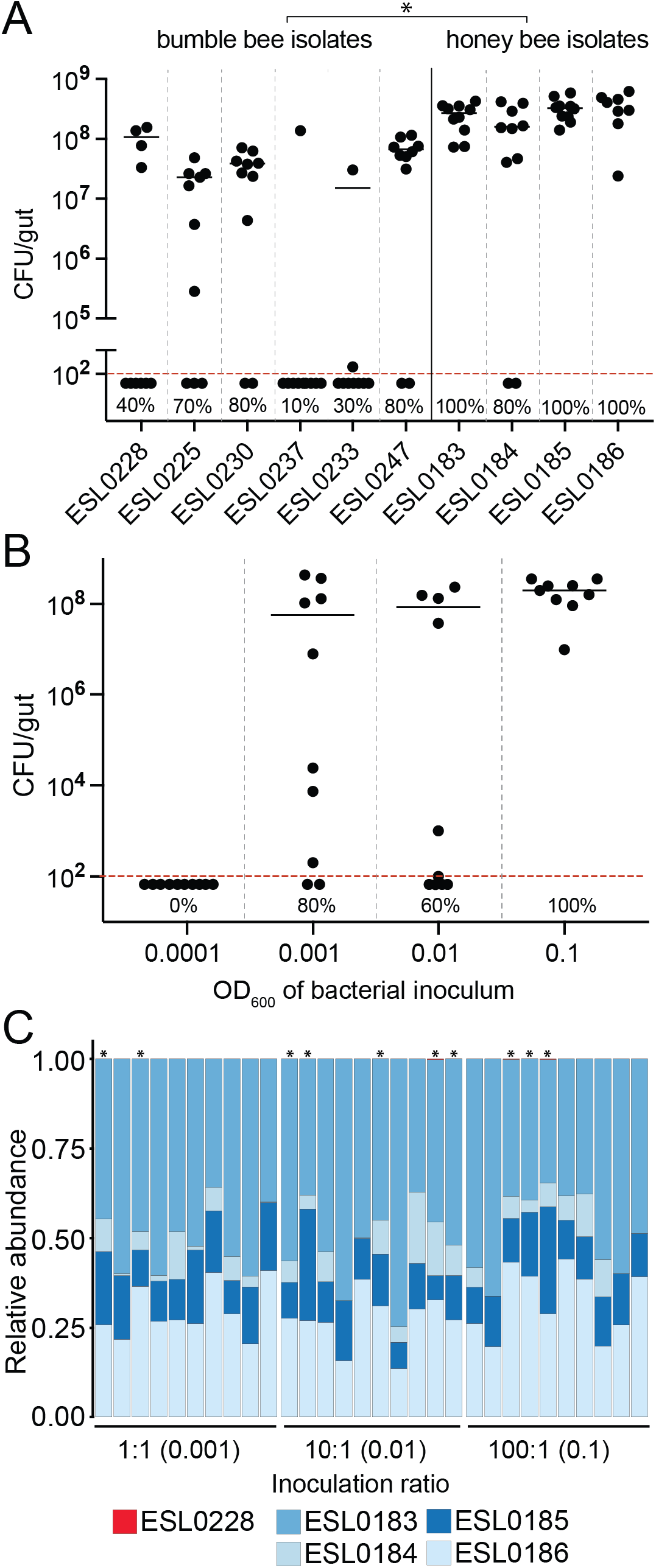
Colonization of microbiota-depleted honey bees with Firm5 strains from bumble bees and honey bees. **(A)** Mono-colonizations of microbiota-depleted honey bees (n=10 per treatment) with six bumble bee strains and four honey bee strains. Each bee was inoculated with 5 μL of an optical density of 0.001. The dashed red line indicates the detection threshold. Data points below the detection limit show bees that had no detectable colonization levels. The percentage of successfully colonized bees is shown. Horizontal lines indicate median. The asterisk above the plot indicates that there is a significant difference between isolates of different host groups in both colonization success (two-sided test of equal proportions, chi=14.1, d.f.=1, p = 0.0002) and colonization efficiency (nested analysis of variance, strains nested within host groups, F=38.1, d.f=1, p = 0.0008). **(B)** Mono-colonizations of microbiota-depleted honey bees with increasing inocula of the bumble bee strain ESL0228. Colony forming units (CFUs) per gut were determined at day 5 post colonization. The graph has the same layout as in panel A. The number of successfully colonized bees and the colonization levels were significantly different across treatments according to a two-sided test of equal proportions (chi-squared = 10.946, df=3, p = 0.0120) and a one way ANOVA (d.f.=3, F=6.646, p = 0.0011). **(C)** Community profiles of microbiota-depleted bees colonized with a community consisting of the bumble bee strain ESL0228 and four honey bee strains (ESL0183, ESL0184, ESL0185, and ESL0186). Three different inoculation ratios of bumble bee strain versus honey bee strains were used. The optical density of the bumble bee strain in the inoculum is given in brackets. Due to the absence or the very low abundance of ESL0228, the red fraction of the graph is not visible. Asterisks indicate samples for which at least a few reads of strain ELS0228 were detected.

To test if the colonization success depends on the number of cells in the inoculum, we colonized microbiota-depleted honey bees with different inocula of the bumble bee Firm5 strain ESL0228 (**Figure S5B**). We chose this particular strain for follow-up experiments, because it had an intermediate colonization success, yet resulted in relatively high bacterial loads compared to the other bumble bee strains. We found a statistically significant difference in the colonization success (two-sided test of equal proportions; chi-squared = 10.946, df=3, p = 0.0120) and the colonization levels (one-way ANOVA, d.f.=3, F=6.646, p = 0.0011) across treatments. With the lowest inoculum, no colonization was obtained at day 5 post colonization (n=10), while with the highest inoculum, all bees were colonized, yielding between 10^6^ – >10^8^ CFUs per gut as in the previous experiment (**Figure 2B**). The relatively high number of bacteria that was needed to achieve a robust colonization suggests that stronger host selection is at play on bumble bee than on honey bee Firm5 strains for gut colonization.

To test for the effect of competitive exclusion between strains, we co-colonized microbiota-depleted bees with the bumble bee Firm5 strain ESL0228 and a mix of four honey bee strains (ESL0183, ESL0184, ESL0185, and ESL0186), each from one of the four divergent sublineages. We kept the number of bacteria in the inoculum constant for the honey bee strains (1:1:1:1), but provided the bumble bee strain at ratios of 1:1, 10:1, or 100:1 relative to the honey bee strains (**Figure S5B**). All bees in the experiment (n=30, n=10 per treatment) were successfully colonized by the Firm5 phylotype and the total numbers of CFUs per gut were in the same range as for the mono-colonizations (10^8^ – 10^9^ CFUs). We used amplicon sequencing of a short fragment of a conserved housekeeping gene to determine the relative abundance of the five Firm5 strains tested in the community (see Methods). This analysis revealed that overall all four honey bee strains successfully colonized and coexisted in the gut, except for strain ESL0184 which was absent from a few samples (**Figure 2C**). In contrast, the bumble bee strain ESL0228 was detected in only a few bees and at very low relative abundance (<0.1%), even when inoculated with a ratio of 100:1.

Collectively, these experiments show that bumble bee strains of the Firm5 phylotype are capable of colonizing the honey bee gut, but consistent colonization can only be achieved with a relatively high inoculum and when honey bee strains are absent. Therefore, we conclude that both host selection and interbacterial competition contribute to the restriction of the realized niche of bumble bee strains.

### Firm5 strains harbor a large gene pool of phylotype-specific functions of which few are conserved

In order to identify genomic characteristics that may contribute to host specialization among Firm5 strains, we carried out a detailed comparative genome analysis. We first determined the distribution of the entire pan genome across the analyzed Firm5 strains. We included 15 divergent outgroup strains in this analysis (i.e. strains not belonging to the Firm5 phylotype, see **Figure 1** and methods) to identify Firm5– specific gene families that could play a role in adaptation to the bee gut environment. In total, 8,248 gene families were identified across the 37 genomes, of which 2,131 gene families were only represented by Firm5 strains (“Firm5-specific”). Of those, 571 and 1,222 gene families were only represented by bumble bee and honey bee strains, respectively, and 338 gene families were represented by members of both hosts (**Figure 3A**).

**Figure 3.**
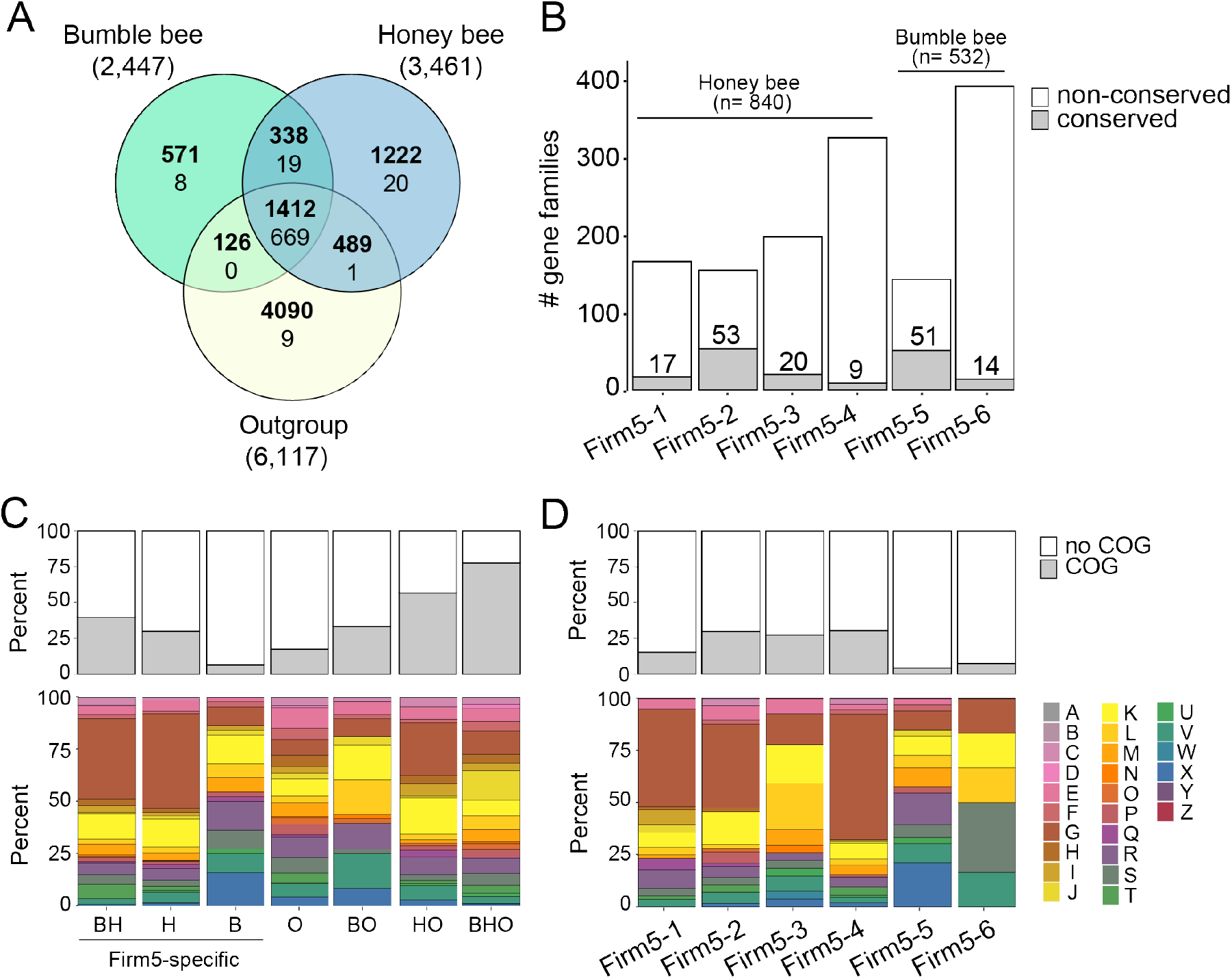
Pan genome analysis of Firm5 strains from honey bees and bumble bees and comparison to outgroup strains (i.e. closely related lactobacilli). **(A)** Venn diagram showing gene family distribution into the three major groups: Firm5 strains from bumble bees, Firm5 strains from honey bees, and outgroup strains. Numbers in bold indicate the total number of gene families (i.e. present in at least one genome of a given group). Numbers in regular font indicate core genome gene families (i.e. present in all genomes of a given group). **(B)** Number of Firm5-specific gene families exclusively present in strains of one sublineage. The fraction of the gene families belonging to the core genome (present in every genome of a given sublineage) and the accessory genome (present in at least one genome of a given sublineage) gene families is indicated by grey and white color, respectively. Numbers above the graph indicate total number of lineage-specific gene families for each host group. **(C)** Upper plot shows number of gene families with COG annotation, and lower plot shows COG category distribution of the annotated gene families for each subset of the Venn diagram in panel A. B, specific to bumble bee strains; H, specific to honey bee strains; O, specific to outgroup strains; BH, shared between honey bee and bumble bee strains; BO, shared between bumble bee and outgroup strains; HO, shared between honey bee and outgroup strains; BHO, shared between all three groups. **(D)** Same as in panel C, but for the sublineage-specific gene families shown in panel B. Complete lists of all gene families and their annotations can be found in **Datasets S1 and S2**. The dominant COG category ‘G’ is shown in dark red and corresponds to ‘Carbohydrate transport and metabolism’. Other COG category abbreviations are given in **Dataset S2**.

Despite this relatively large gene pool of Firm5-specific functions, few gene families were core gene families (defined as gene families shared by all members of a group) (**Figure 3A**). Among the 19 Firm5-specific core gene families, we found an ABC transporter system for branched chain amino acids and two putative adhesin genes (DUF4097). Amino acid transporter genes were also present among the 20 honey bee-specific core gene families, whereas the 8 bumble bee-specific core gene families were annotated as either hypothetical proteins or transcriptional regulators, providing few clues about their functional roles for host adaptation (Dataset S1). Altogether, this analysis revealed very few phylotype-or host-specific gene functions as potential candidates for general determinants of host adaptation across the analyzed Firm5 strains.

### Firm5-specific gene content is restricted to sublineages

As strains from the same host can belong to divergent sublineages, it is possible that sublineage-specific gene functions are involved in host specialization, e.g. by adaptation to different metabolic niches within the gut. Such genes could also explain the ability of the four honey bee sublineages of Firm5 to coexist within bees (Ellegaard & Engel 2019).

Indeed, we found that a relatively large fraction of the Firm5-specific gene families only contained members of a single sublineage (840 of 1,222 for honey bee strains, 532 of 571 for bumble bee strains). However, as for the previous analysis, sublineage-specific core gene families represented a minor fraction (**Figure 3B**). In the honey bee sublineage Firm5-2 *(L. helsingborgensis)* 53 gene families were present in all three strains (i.e. 34% of the sublineage-specific gene content), including several sugar transporter genes and a genomic island for the breakdown of rhamnogalacturonan, a major polysaccharide of pectin (**Dataset S1**). This genomic island was also found in a recent metagenomic study to correlate in abundance with core genes of this sublineage (Ellegaard & Engel 2019), suggesting that rhamnogalacturonan utilization is a conserved function of strains belonging to Firm5-2 *(L. helsingborgensis).* For the other three honey bee sublineages, only 9-20 gene families (0.3%-1.1%) represented sublineage-specific core gene content (**Figure 3B**). In sublineage Firm5-3 *(L. melliventris),* functions for rhamnose utilization were present in all four strains, whereas the annotations of the gene families in the other two sublineages provided little functional insights (**Dataset S1**). The same was the case for the two sublineage-specific core gene families found in bumble bees (9 and 59 gene families, **Figure 3B**), most of which were annotated as hypothetical proteins (**Dataset S1**). Overall, these results indicate a high degree of gene content variability among strains from the same host, also within sub-lineages.

### Honey bee strains harbor a larger diversity of carbohydrate-related functions than bumble bee strains

The high degree of gene content plasticity within the Firm5 phylotype prompted us to look at the functional composition of the entire Firm5-specific gene pool. This analysis revealed marked differences between honey bee and bumble bee strains with respect to carbohydrate-related functions. While ‘Carbohydrate transport and metabolism’ (COG category ‘G’) was by far the most dominant COG category among the gene families specific to the honey bee strains (184 gene families, 50% of those with COG annotation), this category was nearly absent among the gene families specific to the bumble bee strains (4 gene families, 10% of those with COG annotation) (**Figure 3C**). In fact, most gene families specific to bumble bee strains had no COG annotation at all. Analysis of the sublineage-specific gene content revealed a similar pattern. For three of the four honey bee sublineages, COG category ‘G’ was the most abundant COG category, while for the two bumble bee sublineages this category was much less prominent (**Figure 3D**).

This pattern was further explored by quantifying the carbohydrate-related functions within individual genomes. Notably, both the relative and total number of genes assigned to COG category ‘G’ was higher for most honey bee strains compared to bumble bee strains or outgroup strains (**Figure 4A, Figure S6**) (chi-squared=29.647, d.f.=2, p = 3.7*10^-7^). However, the Firm5-1 sublineage represented an exception to this pattern. All three strains of this sublineage encoded fewer COG category ‘G’ genes than other honey bee strains. A large proportion of the genes assigned to COG category ‘G’ encoded phosphotransferase systems (PTSs), i.e. transporters involved in sugar utilization. Correspondingly, these gene families showed a similar distribution as the COG category ‘G’ genes across the Firm5 strains, with most honey bee strains harboring a much larger number of PTS genes than bumble bee strains (**Figure 4B**) (chi-squared = 31.057, d.f. = 2, p = 1.8*10^-7^).

**Figure 4.**
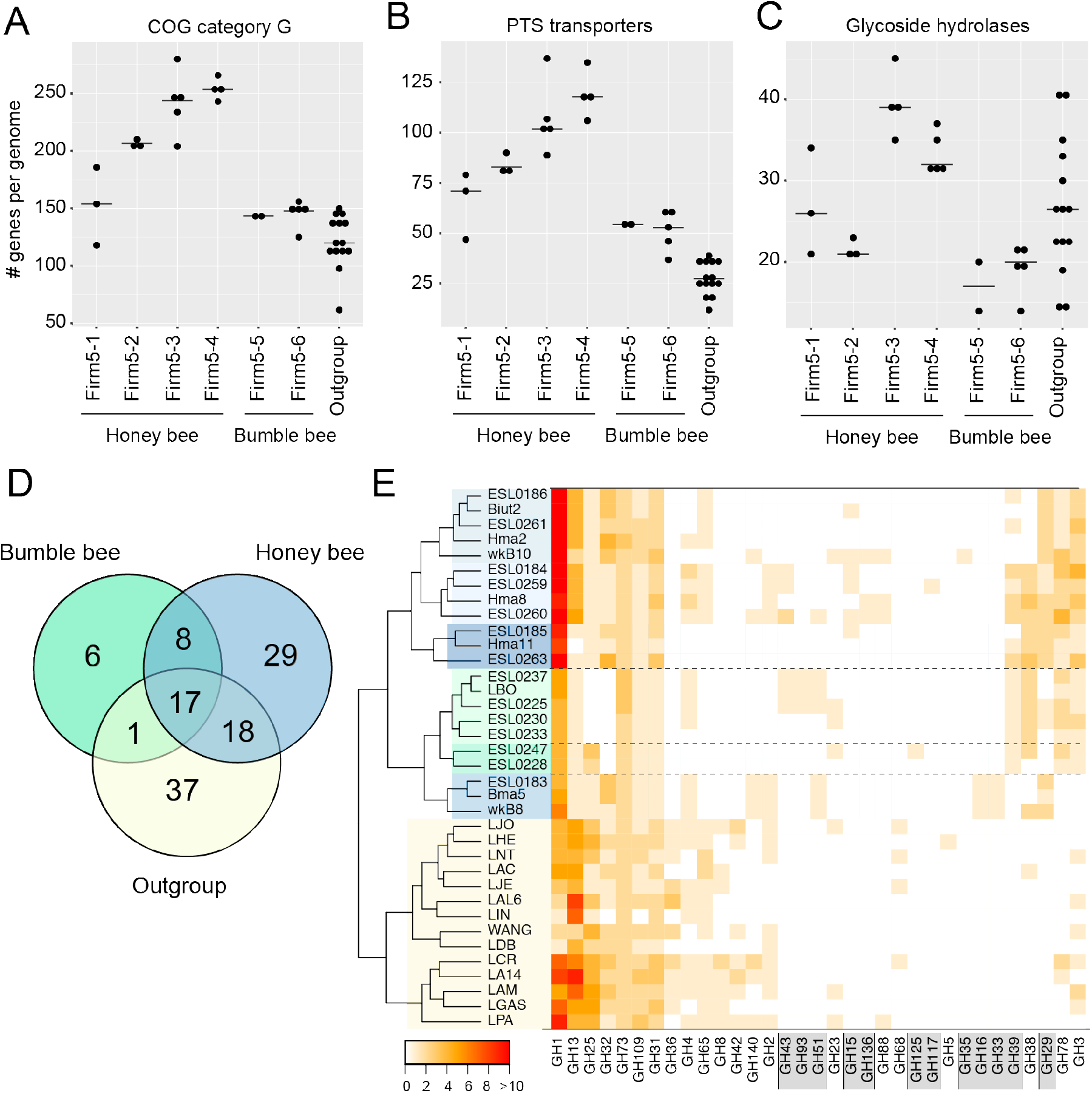
Distribution of carbohydrate-related gene families across strains of the Firm5 phylotype. **(A)** Total number of COG category ‘G’ gene families per genome per sublineage. **(B)** Total number of PTS (Phosphotransferase system) gene families per genome per sublineage. **(C)** Total number of glycoside hydrolase gene families per genome per sublineage. In all three panels, the genomes of the outgroup strains were included as a reference. **(D)** Venn diagram of glycoside hydrolase gene family distribution into the three major phylogenetic groups: Firm5 strains isolated from bumble bees, Firm5 strains isolated from honey bees, and outgroup strains. **(E)** Heatmap showing the distribution of the identified glycoside hydrolase (GH) families across the analyzed genomes. The dendrogram on the left shows a hierarchical clustering based on glycoside hydrolase distribution. Strains are colored according to the major groups (green, bumble bee strains; blue, honey bee strains; yellow, outgroup) and sublineage (color tones). GH families specific to the Firm5 phylotype are indicated by grey boxes.

Within genomes, PTS transport systems are often co-localized with glycoside hydrolases (GHs), which mediate the cleavage of sugar residues from polysaccharides or other glycosylated compounds. To assess if bee gut bacteria harbor a specific arsenal of these sugar-cleaving enzymes, we identified all GH genes in the analyzed genomes. As for COG category ‘G’ and PTS transporters, we found a larger number of GH genes for honey bee strains compared to bumble bee strains (**Figure 4C, Dataset S3**)(chi-squared=11.458, d.f.=2, p = 0.0033). Some honey bee strains harbored twice as many GH genes than bumble bee strains. However, there was remarkable variation in the number of GH genes among the honey bee strains, both within and across sublineages. Specifically, all strains of sublineages Firm5-3 and Firm5-4 harbored a relatively high number of GH genes, while strains of sublineage Firm5-1 varied substantially in the number of GH genes, and those of sublineage Firm5-2 were consistently low.

The identified GH genes belonged to 79 different gene families (**Figure 4D, Dataset S3**), of which 43 were specific to the Firm5 phylotype. Most of these (67%) were only detected among honey bee strains, 19% were shared, and only 14% were specific to bumble bee strains. Moreover, honey bee strains also shared more GH gene families with the outgroup strains than bumble bee strains (18 vs 1 gene families).

While the substrate specificity of GH gene families cannot be unambiguously inferred from sequence data, many of the Firm5-specific GH gene families included glucosidases, fucosidases, mannosidases, xylosidases, and arabinofuranosidases (e.g. GH29, GH38, GH39, GH43, and GH51), as based on the CAZY (Carbohydrate-Active enZYmes) database classification (**Figure 4E**). A similarity search against the publicly available non-redundant database NCBI nr (NCBI Resource Coordinators 2018) revealed that many of the Firm5-specific GH families, especially those exclusively present among the honey bee strains, have best hits to other taxonomic groups than lactobacilliales (**Figure S7**). While these gene families may have been acquired by horizontal gene transfer or secondarily lost in other lactobacillus, their limited distribution among lactobacilliales suggests specific functions in the bee gut environment.

Overall, the analysis of the carbohydrate-related gene content shows that Firm-5 strains from honey bees harbor a larger diversity of PTS transporters and glycoside hydrolases than bumble bee strains. However, differences in the type and abundance of these functions between strains and sublineages suggest that honey bee strains have diversified in their ability to utilize different sugar resources.

### Firm5 strains from bumble bees harbor class II bacteriocins, Firm5 strains from honey bees do not

Most gene families specific to the bumble bee strains were annotated as hypothetical proteins (**Figure 3C and D**), providing no insights about the possible genetic basis of adaptation to the bumble bee gut environment. However, we found several short open reading frames encoding putative class II bacteriocins. Bacteriocins are small peptide toxins that act against closely related bacterial strains (Cotter *et al.* 2003). In the case of class II bacteriocins, an ABC-like transporter usually facilitates toxin secretion, and a dedicated immunity protein provides self-protection. Except for strain ESL0228, all bumble bee strains harbored at least one class II bacteriocin gene with homology to lactococcin 972, described to inhibit septum formation (Martínez *et al.* 2000). Consistent with the genetic organization of lactococcin loci in other species (Letzel *et al.* 2014), putative immunity proteins and ABC transporter genes were encoded downstream of the bacteriocin gene (**Figure 5**). We identified four distinct genomic regions with this genetic organization. All four regions exhibited a high degree of genomic plasticity, with many non-conserved open reading frames close by (**Figure 5 and Figure S8**). Each bacteriocin locus was specific to one of the two bumble bee sublineages and only present in a subset of the analyzed strains. In sublineage Firm5-5, one region encoded two adjacent bacteriocin loci, and in several instances one of the immunity protein or toxin genes was pseudogenized (**Figure 5**). Strikingly, none of these genomic regions were present in the analyzed honey bee strains, suggesting that this genetic feature is specific to bumble bee strains. However, we found homologs of genes for helveticin-J in honey bee strains, another protein with known bactericidal activity against related bacteria. This gene family was conserved in all strains of Firm5 as well as in some of the outgroup strains (**Figure S9**).

**Figure 5.**
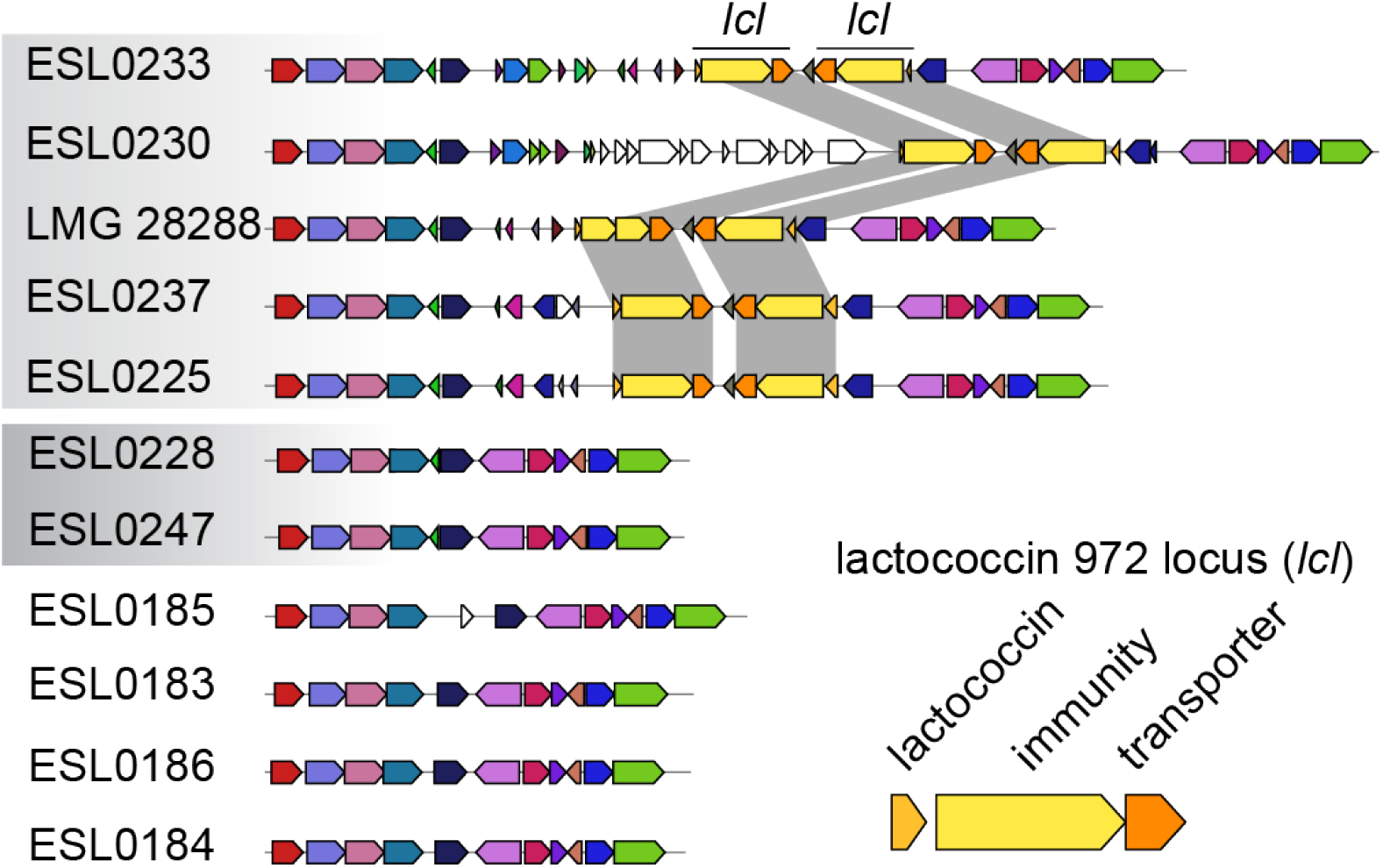
Genomic region encoding class II bacteriocins in Firm5 strains of bumble bee strains. Genomic regions encoding bacteriocin genes were identified and visualized with MultiGeneBlast v1.1.14 (Medema *et al.* 2013). Arrows present genes and same color indicates homology. A black line indicates the lactococcin 972 locus *(lcI)* and vertical grey blocks connect the homologous genes in other strains. An enlarged version of the three genes of the *lcl* locus with annotation is shown in the lower right. Grey shading over strain names indicates two sublineages of bumble bee strains; the four honey bees strains are representatives of the four sublineages. Other genomic regions encoding bacteriocins genes are given in Figure S8.

In summary, while the two bumble bee sublineages of Firm5 harbored a large pool of host-specific gene families, bacteriocins were the only conserved genes with annotated functions, and thus the only identified candidates to play a role for niche specialization in the present state of our knowledge.

## Discussion

In this study, we combined honey bee colonization experiments with comparative genomics to investigate host specialization of Lactobacillus Firm5, a dominant gut symbiont of social bees. Our results show that strains isolated from honey bees and bumble bees belong to separate, highly divergent sublineages of the Firm5 phylotype, which parallels phylogenetic analysis of other bee gut symbionts (Kwong *et al.* 2014; Zheng *et al.* 2016; Kwong *et al.* 2017; Steele *et al.* 2017).

Interestingly, all tested Firm5 strains from bumble bees were able to colonize the gut of microbiota-depleted honey bees, indicating that the divergent evolution of Firm5 strains from different bee species has not resulted in strict host specialization. However, the percentage of successfully colonized bees as well as the number of bacterial cells per gut were both lower for bumble bee strains compared to honey bee strains. Only by increasing the number of bacterial cells in the inoculum by 100-fold were we able to achieve reliable colonization, which suggests strong negative selection of bumble bee strains during passage through the honey bee gut, possibly due to the lack of host-specific adaptation.

However, we currently do not know whether, inversely, bumble bee strains would perform better, and honey bee strains worse, in microbiota-depleted bumble bees, which would provide further evidence for host-specific adaptation. Nevertheless, the fact that strains from both hosts can colonize the honey bee gut extends the fundamental niche of bumble bee strains to other bee genera and shows a partial fundamental niche overlap amongst Firm5 strains. This is in agreement with a previous study showing that selected bacteria from diverse environments, including zebrafish or termite gut, can establish in the gut of germ-free mice (Seedorf *et al.* 2014). Moreover, the gut symbionts *S. alvi* (social bee gut) and *L. reuteri* (vertebrate gut) – for which host specialization has been experimentally demonstrated – are both able to colonize non-native hosts, although at much lower levels than native hosts (Frese *et al.* 2013, Kwong *et al.* 2014).

Niche overlap can result in either niche partitioning, a differential utilization of resources by the organisms involved, or competitive exclusion, where the best competitor drives the other to extinction in the community. (Macarthur & Levins 2015). Although the genomic differences in carbohydrate utilization suggests at least partial niche partitioning between strains that would allow coexistence in the honey bee gut, the bumble bee strain did not establish in any of the tested honey bees when co-inoculated with honey bee strains, even when inoculated with up to 100x more bacterial cells than the four honey bee strains. This clearly shows that the tested bumble bee strain is competitively excluded by the honey bee strains in the honey bee gut. Similar results were obtained for *S. alvi*, when a non-native strain was challenged with a native competitor for gut colonization (Kwong *et al.* 2014).

Honey bees live in large colonies and engage in frequent social interactions. This results in constant exposure to bacteria from nestmates, thereby providing few opportunities for bacteria from non-native hosts to establish in the gut of young worker bees during community assembly. However, even when given the ecological opportunity for gut colonization (as in our colonization experiments), bumble bee strains seem not be able to reliably colonize the gut. Hence we conclude that the competitive disadvantage relative to honey bee strains as well as suboptimal host adaptation both contribute to the exclusion of bumble bee Firm5 strains from the honey bee gut in natural populations.

The bacteria-mediated exclusion of the bumble bee Firm5 strains from the honey bee gut could arise via direct antagonistic interactions between bacteria (e.g. via bacterial toxins), or from resource competition. We identified a number of genes encoding bacteriocins, which are known to mediate interbacterial killing (Kommineni *et al.* 2015). These genes were either shared by strains isolated from both hosts, or they were specific to the bumble bee strains. However, it is notable that the strain selected for the competition experiments, ESL0228, was the only bumble bee strain lacking bacteriocin gene homologs. We can thus not exclude at this point that bumble bee strains carrying bacteriocin genes would be more competitive in the honey bee gut. Vice versa, we did not identify any toxin genes specific to the honey bee strains, which could mediate possible antagonistic effects towards bumble bee strains and hence hinder their colonization in the gut.

Biofilm formation at the host epithelium has been shown to be a crucial factor for the colonization success and competitiveness of the murine gut symbiont *L. reuteri* (Frese *et al.* 2013; Duar *et al.* 2017). The honey bee gut symbiont *S. alvi* also colonizes the epithelial surface and forms biofilm-like structures, making it conceivable that competition for adherence is also a critical factor for colonization in the bee gut (Kwong & Moran 2016). However, bacteria of the Firm5 phylotype do not seem to attach to the host epithelium, as shown by fluorescence in situ hybridization experiments, but rather colonize the gut lumen in the rectum (Martinson *et al.* 2012), where competition for space seems less likely to be a predominant limiting factor. Moreover, our genomic analysis did not identify genes involved in host interaction or adherence to be specific to strains from one of the two host groups.

Instead our genomic analysis showed that the strains differ in terms of the quantity and diversity of metabolic functions. Although we found relatively few honey bee-specific core gene families, the genomes of honey bee strains consistently harbored a larger arsenal of genes related to carbohydrate metabolism and transport compared to bumble bee and outgroup strains. This suggests that honey bee strains have a greater capacity to utilize diet-derived carbohydrates, which may give them a growth advantage over bumble bee strains in the bee gut. A similar trend has also been observed for strains of the gut symbiont *G. apicola* (Kwong *et al.* 2014).

The predominant energy metabolism of the Firm5 phylotype is predicted to be fermentation of dietary carbohydrates, which is not surprising given that the diet of social bees (pollen and nectar) is rich in simple sugars, polysaccharides (pectin, hemicellulose and cellulose), and other glycosylated compounds (e.g. flavonoids) (Engel *et al.* 2012; Ellegaard *et al.* 2015; Kešnerová *et al.* 2017). However, bumble bees and honey bees have a similar dietary regime, as both eat nectar and pollen. Hence, the reason for why bumble bee strains harbor significantly fewer carbohydrate-related functions is currently unclear. Interestingly, almost none of the carbohydrate-related gene families specific to honey bee strains of the Firm5 phylotype were conserved across the analyzed genomes, suggesting that the genetic basis of host adaptation in regard of carbohydrate metabolism differs between strains. Moreover, although a large proportion of the carbohydrate-related gene content was specifically associated with one of the four sublineages of honey bee Firm5 strains, only a few of these functions were conserved within sublineages (e.g rhamnogalacturonan and rhamnose utilization in Firm5-2 and Firm5-3, respectively), and the number of carbohydrate-related functions varied markedly among strains of some sublineages. Taken together, these results suggest that metabolic functions are also more frequently gained and lost in honey bee strains compared to bumble bee strains.

This could possibly be related to known differences in the life cycle of bumble bees and honey bees. Honey bees maintain perennial colonies of large population sizes (ca. 20-50,000), while bumble bees build smaller colonies (typically < 500 individuals) from a single overwintering queen every year. This represents a population bottleneck for the bacterial community in the gut of bumble bees, since only bacteria colonizing the queen are expected to be transferred to colony members in the following season. Likewise, smaller colonies would reduce the effective population size, unless migration across colonies is frequent. If so, genetic drift would lead to gene loss and decrease the selective pressure imposed by related bacteria. It would also slow the acquisition of novel gene functions allowing bacteria to utilize diverse carbohydrates. Strikingly, in the host-specialized vertebrate gut symbiont *L. reuteri*, it was also speculated that genomic differences in genome size and pan genome diversity may be due to differences in population bottlenecks across hosts (Frese *et al.* 20131).

In conclusion, our study advances the understanding of host specialization of gut symbionts. While previous studies on *L. reuteri* have shown that host interaction, and specifically colonization of the gut surface, determine host specificity, we provide evidence for metabolic flexibility that may facilitate adaptation to the host diet and hence the competitive exclusion of non-adapted strains. As specific dietary preferences are common among animals, similar processes may also be a determining factor of host specialization among other gut symbionts.

## Materials and methods

### Bee sampling, bacterial culturing and DNA isolation

Bumble bees were collected from flowers in different locations in Western Switzerland as indicated in **Table S1**. Honey bees were sampled from two healthy looking colonies in the same region located at the University of Lausanne. Within 6h after sampling, bees were immobilized on ice and the entire gut was dissected with sterile scissors and forceps. Each gut tissue was individually placed into a screw cap tube containing 1ml 1x PBS and glass beads (0.75-1mm, SIGMA) and homogenized with a bead-beater (FastPrep-24 5G, MP Biomedicals) for 30s at speed 6.0. Serial dilutions of the gut homogenates were plated on MRS agar and incubated at 34°C in an anaerobic chamber (Coy laboratories, MI, USA) containing a gas mix of 8% H_2_, 20% CO_2_ and 72% N_2_. After 3-5 days of incubation, single colonies were picked, restreaked on fresh MRS agar and incubated for another 2-3 days. Then, a small fraction of each restreaked bacterial colony was resuspended in lysis buffer (10 mM Tris-HCl, 1 mM EDTA, 0.1% Triton, pH 8, 2 mg/ml lysozyme and 1mg/ml proteinase K) and incubated in a thermocycler (10 min 37°C, 20 min 55°C, 10 min 95°C). Subsequently, a standard PCR with universal bacterial primers (5’-AGR GTT YGA TYM TGG CTC AG-3’, 5’-CCG TCA ATT CMT TTR AGT TT-3’) was performed on 1 μl of the bacterial lysate and the resulting PCR products were sent for Sanger sequencing. Sequencing reads were inspected with Geneious v6 (Biomatters Limited) and compared to the NCBI nr database (NCBI Resource Coordinators 2018) using BLASTN. Isolates identified to have high similarity (i.e. >95% sequence identity) to honey bee strains of the Firm-5 phylotype were stocked in MRS broth containing 25% glycerol at −80°C. Genomic DNA was isolated from fresh bacterial cultures of the strains of interest using the GenElute Bacterial Genomic DNA Kit (SIGMA) according to manufacturers instructions. Bumble bees were genotyped based on the COI gene by performing a PCR on DNA extracted from the carcass with primers LepF1 and LepR1 (Hebert *et al.* 2004), sending the PCR product for Sanger sequencing, and searching the resulting sequence read by BLASTN against the NCBI nr database.

### Genome sequencing, assembly and annotation

Genome sequencing libraries were prepared with the TruSeq DNA kit and sequenced on the MiSeq platform (Illumina) using the paired-end 2×2 50-bp protocol at the Genomic Technology facility (GTF) of the University of Lausanne. The preliminary genome sequence analysis was carried out in the framework of the student course ‘Sequence-a-genome (SAGE)’ at the University of Lausanne in 2016-2017. In short, the resulting sequence reads were quality-trimmed with trimmomatic v0.33 (Bolger *et al.* 2014) to remove adapter sequences and low quality reads using the following parameters: ILLUMINACLIP:TruSeq3-PE.fa:3:25:6 LEADING:9 TRAILING:9 SLIDINGWINDOW:4:15 MINLEN:60. The quality-trimmed reads were assembled with SPAdes v.3.7.1 (Bankevich *et al.* 2012), using the “--careful” flag and multiple k-mer sizes (-k 21,33,55,77,99,127). Small contigs (less than 500 bp) and contigs with low kmer coverage (less than 5) were removed from the assemblies, resulting in 11–24 contigs per assembly. The contigs of each assembly were re-ordered according to the complete genome of the honey bee strain ESL0183 using MAUVE v2.4 (Rissman *et al.* 2009). The origin of replication was set to the first base of the *dnaA* gene, which coincided with the sign change of the GC skew. The ordered assemblies were checked by re-mapping the quality-trimmed reads (**Figure S2**). Except for a few prophage regions that showed increased read coverage, no inconsistencies in terms of read coverage or GC skew were revealed suggesting that the overall order of the contigs was correct. The median read coverage of the sequenced genomes ranged between 135x-223x (**Figure S2**). The genomes were annotated using the ‘Integrated Microbial Genomes and Microbiomes’ (IMG/mer) system (Markowitz *et al.* 2014).

### Inference of a genome-wide phylogeny

Gene families, i.e. groups of homologous genes, were determined using OrthoMCL (Li *et al.* 2003) between all publicly available and newly sequenced genomes of the Firm5 phylotype as well as a set of outgroup genomes of other lactobacilli strains. The outgroup strains were selected based on their phylogenetic relatedness with the Firm5 phylotype using a previously published phylogeny of the entire genus *Lactobacillus* (Zheng *et al.* 2015). Based on this analysis, we included the genomes of 15 closely related outgroup strains that belong to the same *Lactobacillus* clade as Firm5 (‘delbrueckii group’) and three more distantly related strains for rooting the phylogeny. All-against-all BLASTP searches were conducted with the proteomes of the selected genomes, and hits with an e-value of ≤10^-5^ and a relative alignment length of >50% of the query and the hit protein lengths were kept for OrthoMCL analysis. All steps of the OrthoMCL pipeline were executed as recommended in the manual and the mcl program was run with the parameters ‘--abc -I 1.5’.

The core genome phylogeny was inferred from 408 single copy orthologs extracted from the OrthoMCL output (i.e. gene families having exactly one representative in every genome in the analysis). The protein sequences of each of these core gene families were aligned with mafft (Katoh *et al.* 2017). Alignment columns represented by less than 50% of all sequences were removed and then the alignments were concatenated. Core genome phylogenies were inferred on the concatenated trimmed alignments using RAxML (Stamatakis 2014) with the PROTCATWAG model and 100 bootstrap replicates.

### Comparison of genome structure, genome divergence, and gene content

To compare and visualize whole genomes we used the R-package genoPlotR (Guy *et al.* 2010). BLASTN comparison files were generated with DoubleACT (www.hpa-bioinfotools.org.uk) using a bit score cutoff of 100. To estimate sequence divergence between genomes, we calculated pairwise average nucleotide identity (ANI) with OrthoANI (Lee *et al.* 2016) using the exectutable ‘OAT_cmd.jar’ with the parameter ‘method ani’.

For analyzing the distribution of gene families across Firm5 sublineages and closely related outgroup strains, we carried out a second OrthoMCL analysis, in which we excluded the three distantly related outgroup strains. To remove redundancy in our database, we also excluded the genomes of five Firm5 isolates that were identical, or almost identical, to other Firm5 strains in the analysis, based on ANI values of >99.99%. This resulted in a total 37 genomes (22 Firm5 genomes and 15 outgroup genomes) that were included in the analysis. BLASTP and OrthoMCL were run with the same parameters as before. Gene family subsets of interest (e.g. families specific to honey bee, bumble bee or outgroup strains) were extracted from the OrthoMCL output file using custom-made Perl scripts. COGs (Cluster of Orthologous Groups) were retrieved from IMG/mer genome annotations (Markowitz *et al.* 2014).

For the detection and visualization of the genomic regions encoding bacteriocin genes, Bagel3 (van Heel *et al.* 2013) and MultiGeneBlast (Medema *et al.* 2013) were used. For the MultiGeneBlast analysis, bacteriocins-encoding genomic regions of strains ESL0233 and ESL0247 served as query sequences for searching a custom-made database composed of all non-redundant Firm5 genomes.

### Identification of glycoside hydrolase gene families

Glycoside hydrolase gene families were identified in all analyzed genomes (excluding the redundant Firm5 strains and the three distant outgroup strains) using the command-line version of dbCAN (Database for automated Carbohydrate-active enzyme Annotation) (Yin *et al.* 2012). In short, we searched each genome against dbCAN using hmmscan implemented in HMMER v3 (Eddy 2009). The output was processed with the parser script ‘hmmscan-parser.sh’, and genes with hits to Hidden Markov Models of glycoside hydrolase families were extracted (for alignments > 80aa an e-value cut-off of < 10^-5^ was used, otherwise an e-value cut-off of <10^−3^ was used, the covered fraction of the HMM had to be > 0.3).

For determining the taxonomic distribution of related genes, we searched one homolog of each glycoside hydrolase gene family against the NCBI *nr* database (NCBI Resource Coordinators 2018) using BLASTP. The taxonomy of the first 50 BLASTP hits (e-value <10^-5^) was extracted at the family level using the Perl script ‘Tax_trace.pl’ and the database files nodes.dmp and names.dmp. The latter two files contain the NCBI taxonomy nodes and names.

### Bee colonization experiments

Newly emerged, microbiota-depleted bees were generated as described in (Emery *et al.* 2017) and colonized within 24-36h after pupal eclosion. To this end, bacterial strains were grown on MRS agar containing 2% fructose and 0.2% L-cysteine-HCl from glycerol stocks for two days in an anaerobic chamber at 34°C. Then, 1-10 colonies were inoculated into 5ml of carbohydrate-free MRS supplemented with 4% fructose, 4% glucose and 1% L-cysteine-HCl and incubated for another 16-18h without shaking. Bacteria were spun down and resuspended in 1xPBS/sugar water (1:1). The optical density (600nm) was adjusted according to the experimental condition (OD=0.0001, 0.001, 0.01, or 0.1) and 5 μl of the final bacterial suspension was fed to each newly emerged bee. Before feeding, the bees were starved for 2-3h. After colonization, bees were given 1 ml of sterilized polyfloral pollen and sugar water ad libitum. Bees were co-housed in groups of 20-40 bees. For the competition experiment, each of the four honey bee strains was adjusted to an optical density (600nm) of 0.001. The bumble bee strain ESL0228 was adjusted to an optical density (600nm) of either 0.001, 0.01, or 0.1. Then equal volumes of the five strains were mixed together and fed to newly emerged bees as described before. As negative control, bees were fed with 5 μl of 1xPBS/sugar water. Dilutions of the bacterial inocula were plated on MRS agar containing 2% fructose and 0.2% L-cysteine-HCl and incubated as described before to determine how many CFUs correspond to a given optical density (see Figure S5).

Ten bees per condition were dissected on day 5 after colonization. The hindgut was separated from the midgut with a sterile scalpel and tweezers, and added to 1 ml or 500 ul of 1x PBS (depending on the experiment). The tissues were homogenized by bead-beating as described before, dilutions plated on MRS agar containing 2% fructose and 0.2% L-cysteine-HCl and the number of CFUs counted two to three days after incubation. For the negative control, bacterial colonies were detected for only one out of 30 bees with a relatively low abundance (10^3^ CFUs per gut). Moreover, the colonies looked different from the colonies of the Firm5 strains and were identified as being *E. coli* and *Staphylococcus aureus* by 16S rRNA gene sequencing.

The relative abundance of the five strains in the competition experiment was analyzed using amplicon sequencing of a 199-bp fragment of a conserved housekeeping gene (COG0266). To this end, a two-step PCR protocol was established. In the first PCR, the 199-bp fragment of COG0266 was amplified from crude cell lysates of gut homogenates with primers 1133 (5’ – CGTACGTAGACGGCCAGTATGCCNGAAATGCCRGARGTTGA – 3’) and 1134 (5’ – GACTGACTGCCTATGACGACTAARCGATAYTTRCCYTCCATRCG) (3’ – 95°C; 25x: 30” – 95°C, 30” – 64°C, 30” – 72°C; 5’ – 72°C). After removing primers with exonuclease and shrimp alkaline phosphatase, barcoded Illumina adapters were added in the second PCR. The resulting PCR products were pooled at equal volumes, gel purified (MinElute Gel Extraction Kit, Qiagen) and loaded on an Illumina MiniSeq instrument in mid-output mode. Reads were demultiplexed and filtered on quality using trimmomatic (LEADING:28 TRAILING:28 SLIDINGWINDOW:4:15 MINLEN:90) (Bolger *et al.* 2014).

Then, each forward and reverse read pair was assembled using PEAR (-m 290 -n 284 -j 4 -q 26 -v 10 -b 33) (Zhang *et al.* 2014). The resulting contigs were assigned to the five strains based on base positions with discriminatory SNP variants with the help of a custom-made Perl script.

### Statistical Analysis

Colonization success (as proportion of bees in trial successfully colonized above detection limit) was compared in all experiments with a two-sided test of equal proportions across groups, and again pairwise between all strains/treatments. Differences in colonization efficiency (as the number of CFUs per gut in those bees that were successfully colonized), were tested with a nested analysis of variance where strains are nested within host groups, or with a one-way analysis of variance for the initial inoculum experiment. Normality was tested with a Shapiro-Wilk test and an inspection of the QQ-plots of both raw data and residuals. P-values in post-hoc tests were adjusted with the Benjamin-Hochberg correction (Benjamini & Hochberg 1995) for multiple comparisons, and were considered significant below 0.05. All tests were performed in R.

Genome length and differences in functional gene number were tested with a Kruskal-Wallis test and a Games-Howell post-hoc test, for COG ‘G’ and PTS categories, and with a Dunn’s post-hoc test for genome length and GH.

## Acknowledgments

We thank the School of Biology of the University of Lausanne for financial support of this project. We would also like to thank Ambrin Farizah Babu, Melvin Bérard, Sarah Berger, Laurent Casini, Joaquim Claivaz, Yassine El Chazli, Jonas Garessus, Nastassia Gobet, Charlotte Griessen, Olivier Gustarini, Karim Hamidi, Dominique Jacques-Vuarambon, Titouan Laessle, Mirjam Mattei, Cyril Matthey-Doret., Jennifer Mayor, Sandrine Pinheiro, Claire Pralong, Virginie Ricci, Shaoline Sheppard, Tatiana Sokoloff, Anthony Sonrel, and Gaëlle Spack who participated as students in the SAGE class 2016/2017 and were involved in a preliminary analysis of the genomic data. **Some of** the computations were performed at the Vital-IT (http://www.vital-it.ch) Center for high-performance computing of the SIB Swiss Institute of Bioinformatics. This research was funded through the School of Biology of the University of Lausanne, the ERC-StG ‘MicroBeeOme’, the Swiss National Science Foundation grant 31003A_179487, and the HFSP Young Investigator grant RGY0077/2016.

## Data accessibility

Genome sequences and short read datasets are available under NCBI Bioproject accession PRJNA392822. Annotations of the Firm5 strains used for this study can be found in IMG/mer. Data analyses including custom scripts and intermediate output files are available on Zenodo Data analyses including custom scripts and intermediate output files are available on Zenodo: https://doi.org/10.5281/zenodo.1010076.

## Supplementary Figures

**Figure S1.**
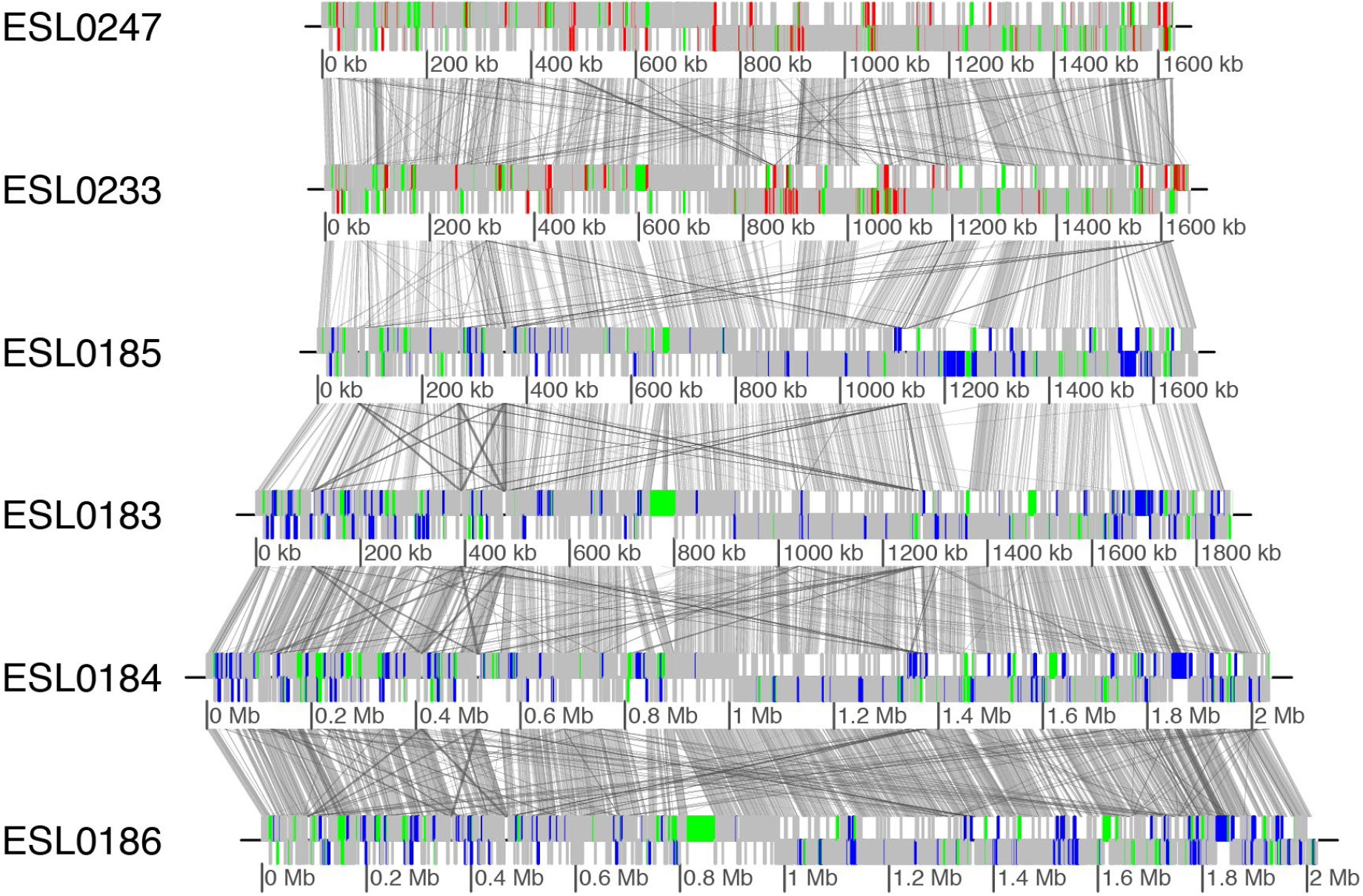
Whole genome alignments of divergent Firm5 strains. ESL0247 and ESL0233 are bumble bee strains. ESL0183-186 are honey bee strains. Vertical grey lines indicate blocks of nucleotide sequence similarity. Different color intensities correspond to different degree of similarity based on BLASTN hits with a bit score of at least 100. Genes in color correspond to Firm5-specific genes relative to the outgroup (blue, genes specific to honey bee strains; red, genes specific to bumble bee strains; green, genes shared between honey bee and bumble bee strains). Other genes are shown in grey.

**Figure S2.**
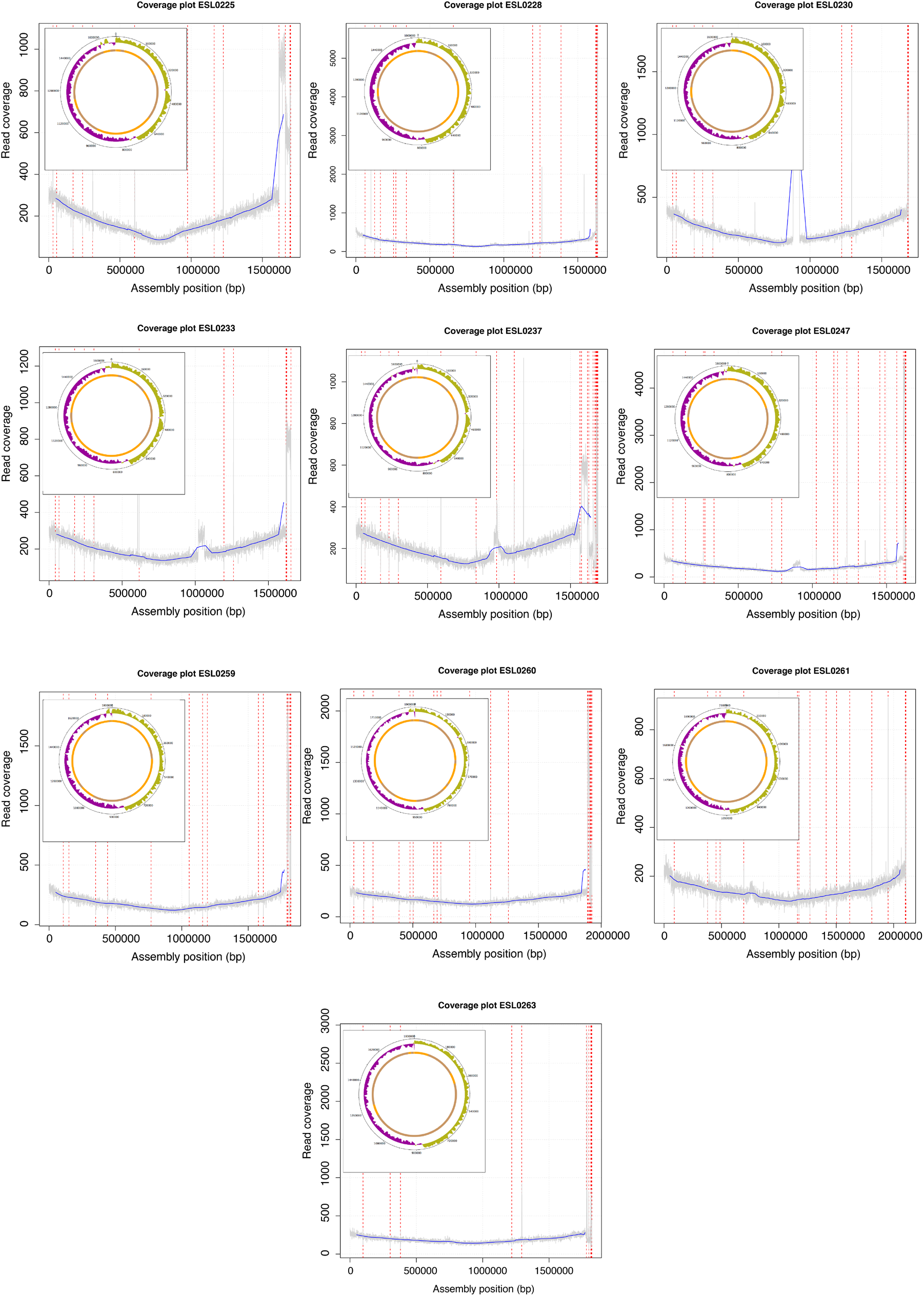
Read coverage and GC skew of the final genome assemblies. Assembly positions are shown on the x-axis for each sequenced strain. y-Axis shows Illumina read coverage for a sliding window of 100 bp. Red dashed lines indicate contig breaks. Contigs were ordered according to the fully sequenced reference strains ESL0183. Small contigs were left at the end of the assembly. Inset shows the circular form of the assembly with the GC skew indicated. We found a higher read coverage at the origin of replication, which is characteristic for replicating bacteria. Moreover, our assemblies showed the typical GC skew of bacterial genomes. Both characteristics indicate that the contigs of the assemblies were correctly assembled and ordered. Notably, regions of extremely high coverage correspond to prophages that apparently were amplified during culturing of some of the strains.

**Figure S3.**
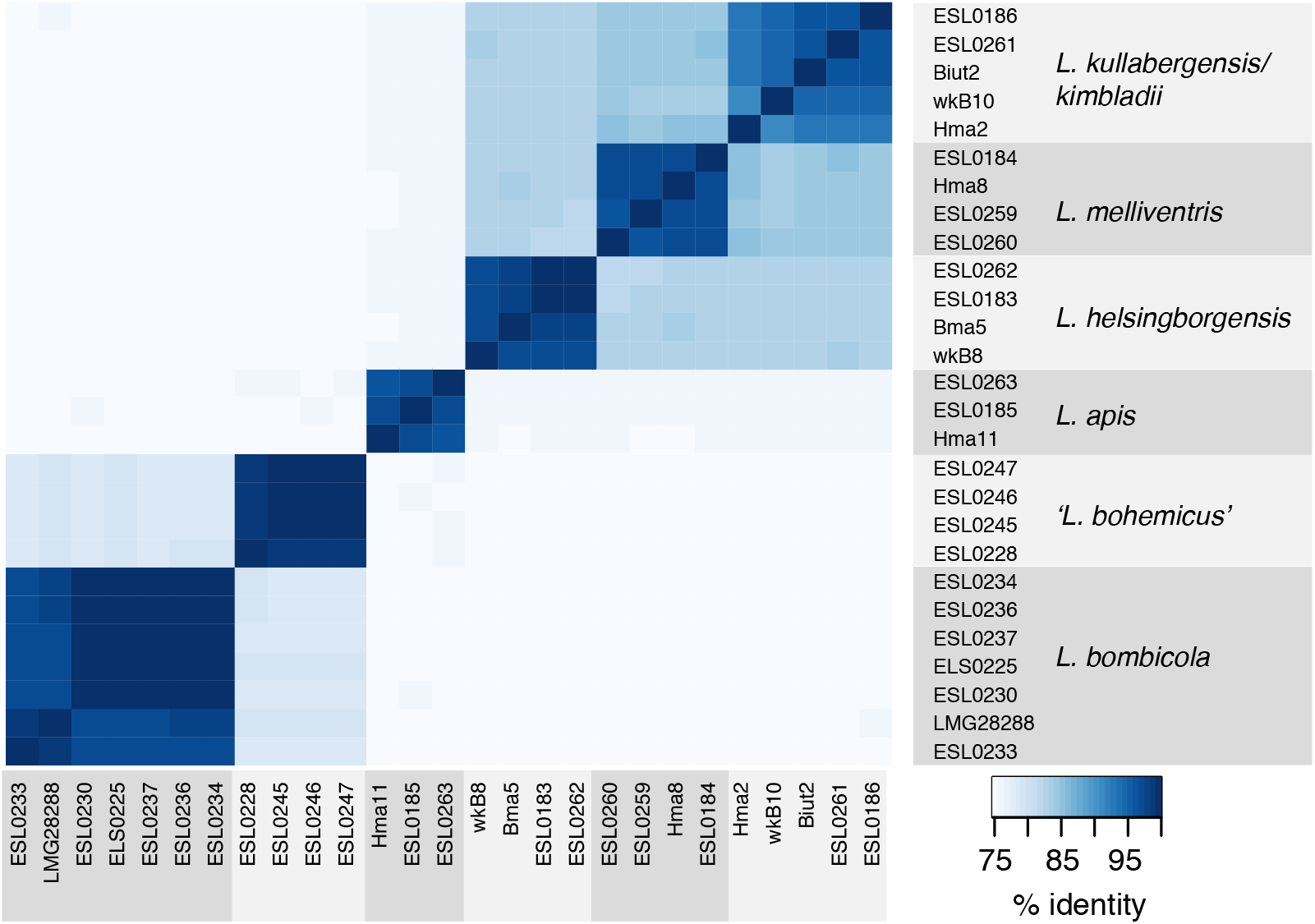
Average nucleotide identity between the analyzed Firm5 genomes. Intensity of heatmap indicates pairwise ANI. White areas correspond to genomes, which were too divergent for ANI calculation. The names of each strain included in the analysis are given next to the plot area (see also **Table S1**). Grey shading indicates the six different sublineages of Firm5. ANI values are given in **Table S3**.

**Figure S4.**
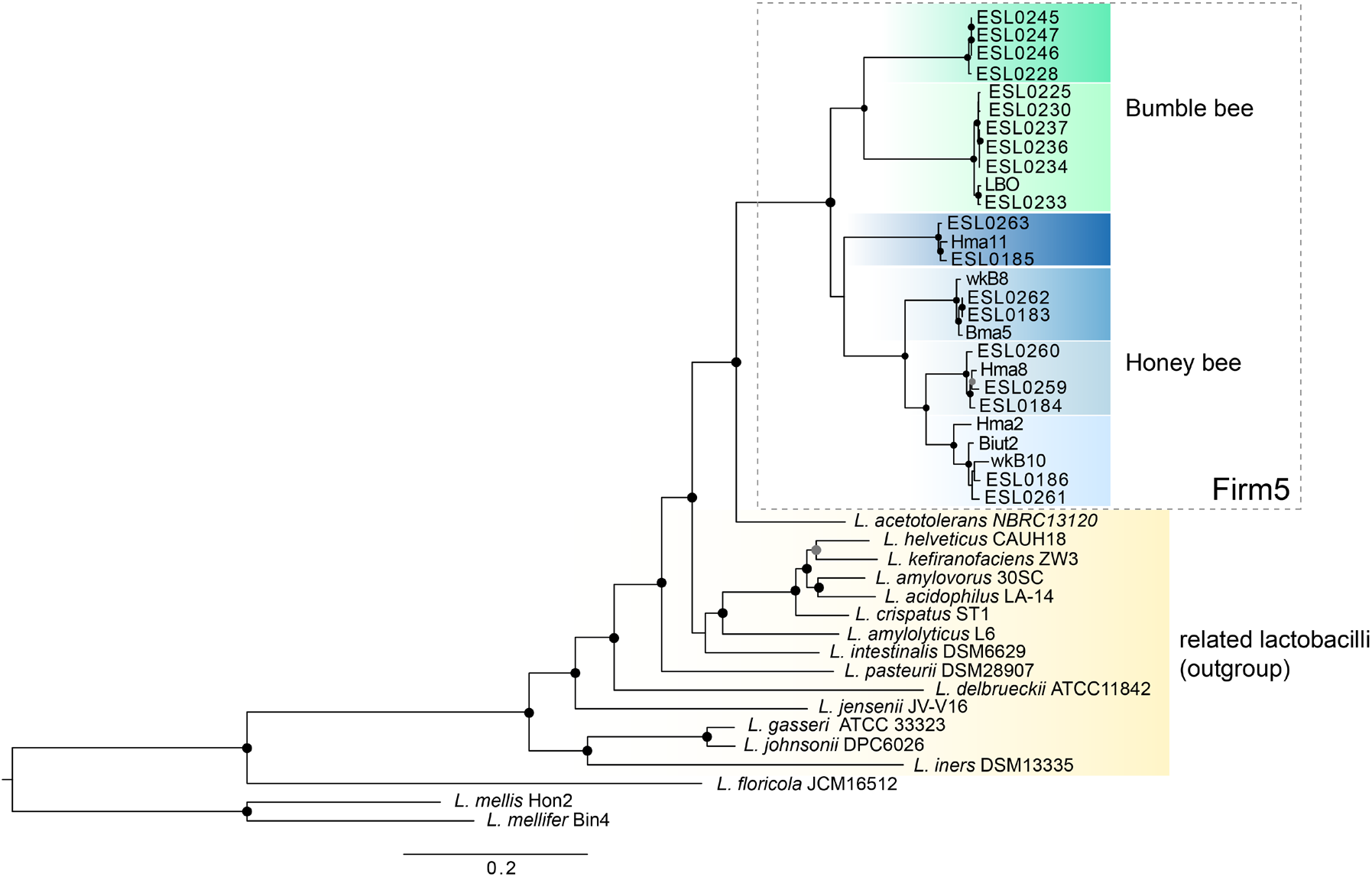
Complete core genome phylogeny of *Lactobacillus* Firm5. The tree was inferred using maximum likelihood on the concatenated protein alignments of 408 single-copy core gene families (i.e. present in all Firm5 strains and the outgroup strains). The two lineages of bumble bee strains and the four lineages of honey bee strains are shown in green and blue color shades, respectively. As outgroup, 15 representative strains of the *L. delbrueckii* group (to which Firm5 belongs to) were included in the analysis (shown in yellow) based on a previously published phylogeny of the entire genus *Lactobacillus* (Zheng *et al.* 2015). In addition, we included three more distantly related strains to root the tree. Noteworthy, these three distantly related lactobacilli were excluded for all subsequent comparative analysis of the Firm5 strains. Filled circles indicate 100 bootstrap support values. The strain designation of each isolate is given. The length of the bar indicates 0.05 amino acid substitutions / sites.

**Figure S5.**
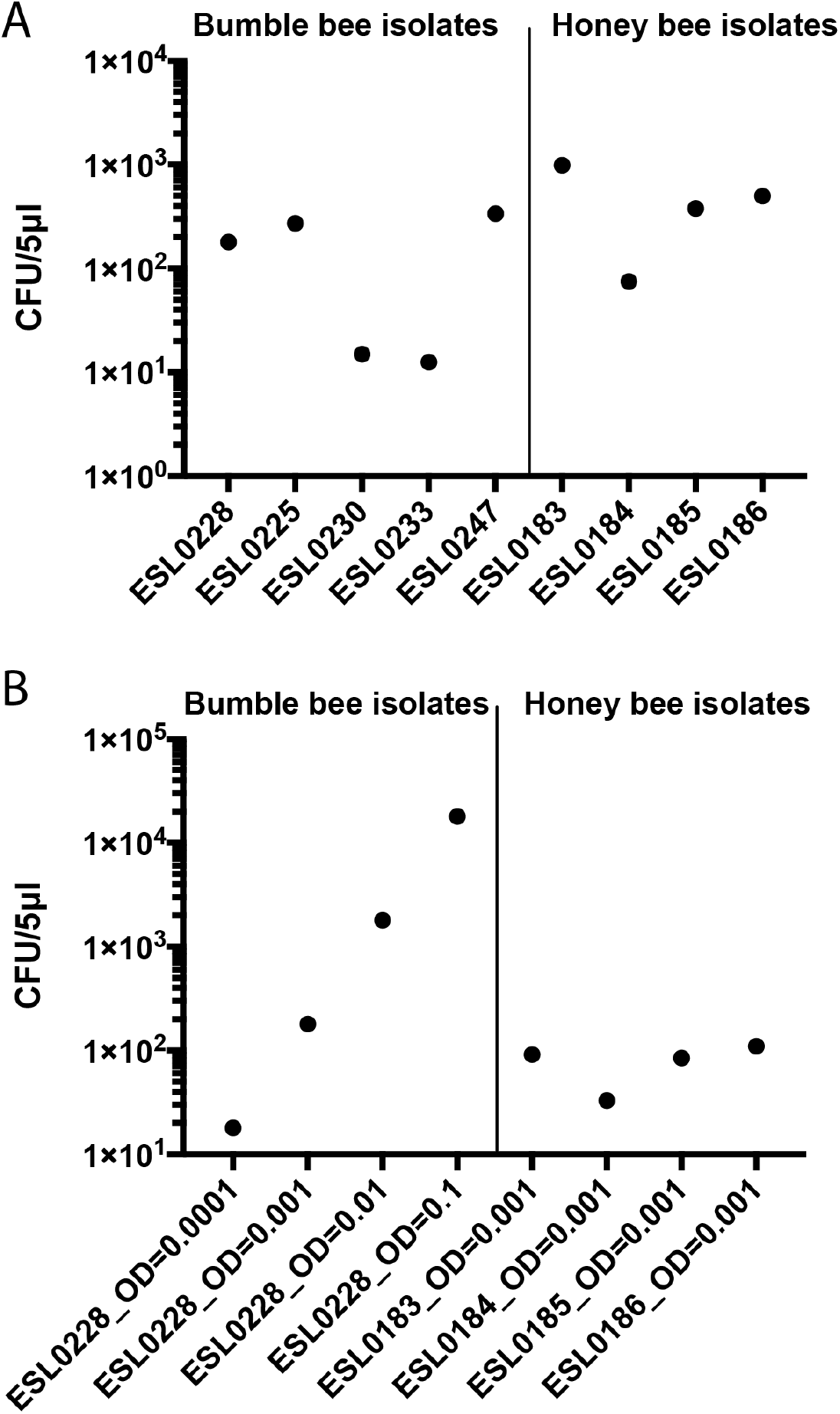
Number of bacterial cell in the inocula used to colonize microbiota-depleted honey bees with Firm5 strains. **(A)** CFUs in the inocula used for the monocolonization experiments with individual strains. CFUs are given per 5μl, as each bee was inoculated with 5 μl of an 0D600 of 0.0001. Despite the adjustment to the same OD600, the amount of live bacteria in each inoculum varied across strains. The inoculum of strain ESL0237 could not be assessed due to a handling mistake during dilution plating. **(B)** CFUs in the incocula used for the colonization experiment with the bumble bee strain ESL0228 (left part) and for the colonization experiment with the five-member community consisting of the bumble bee strain ES0228 (left panel, OD=0.001, 0.01, and 0.1) and the four different honey bee strains (right panel).

**Figure S6.**
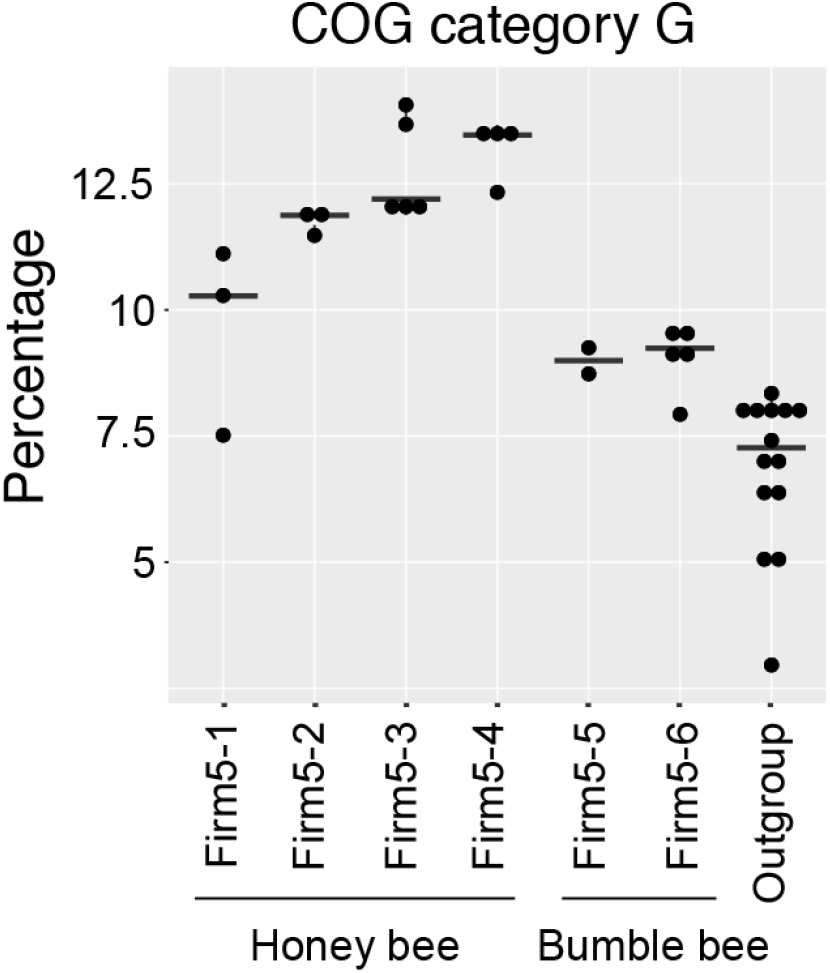
Percentage of gene families annotated as COG category ‘G’ per genome per sublineage. Same data as in Figure 4A, but expressed in relative numbers (percentage of all gene families per genome).

**Figure S7.**
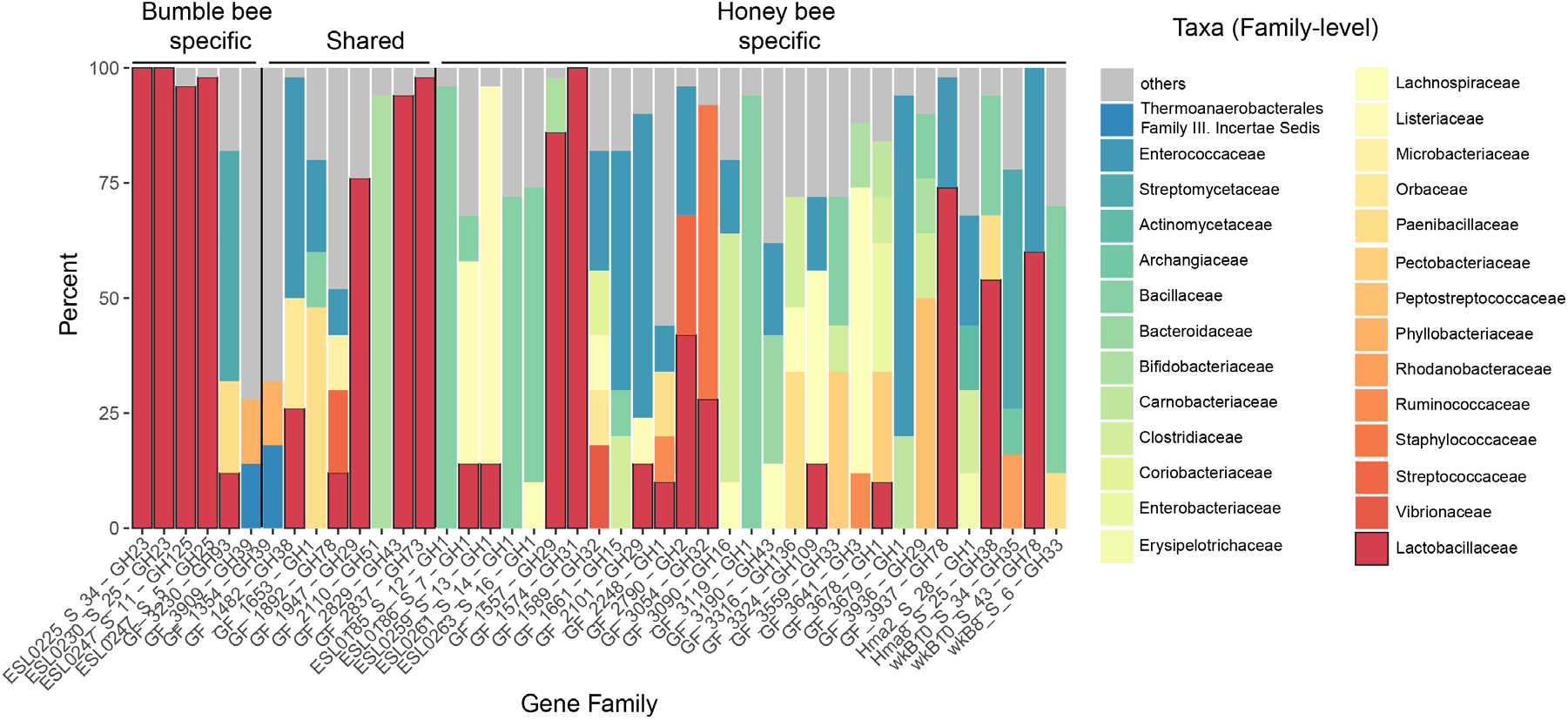
BlastP hit distribution of glycoside hydrolase (GH) gene families specific to Firm5. A representative protein sequence of each gene family was blasted against the NCBI nr database (NCBI Resource Coordinators 2018). The distribution of the taxonomic classification of the first 50 Blast hits is shown at the family level. The family of lactobacilliales is shown in red with black outlines. For each gene family, the gene family identifier and the glycoside hydrolase enzyme family (GHxx) are given.

**Figure S8.**
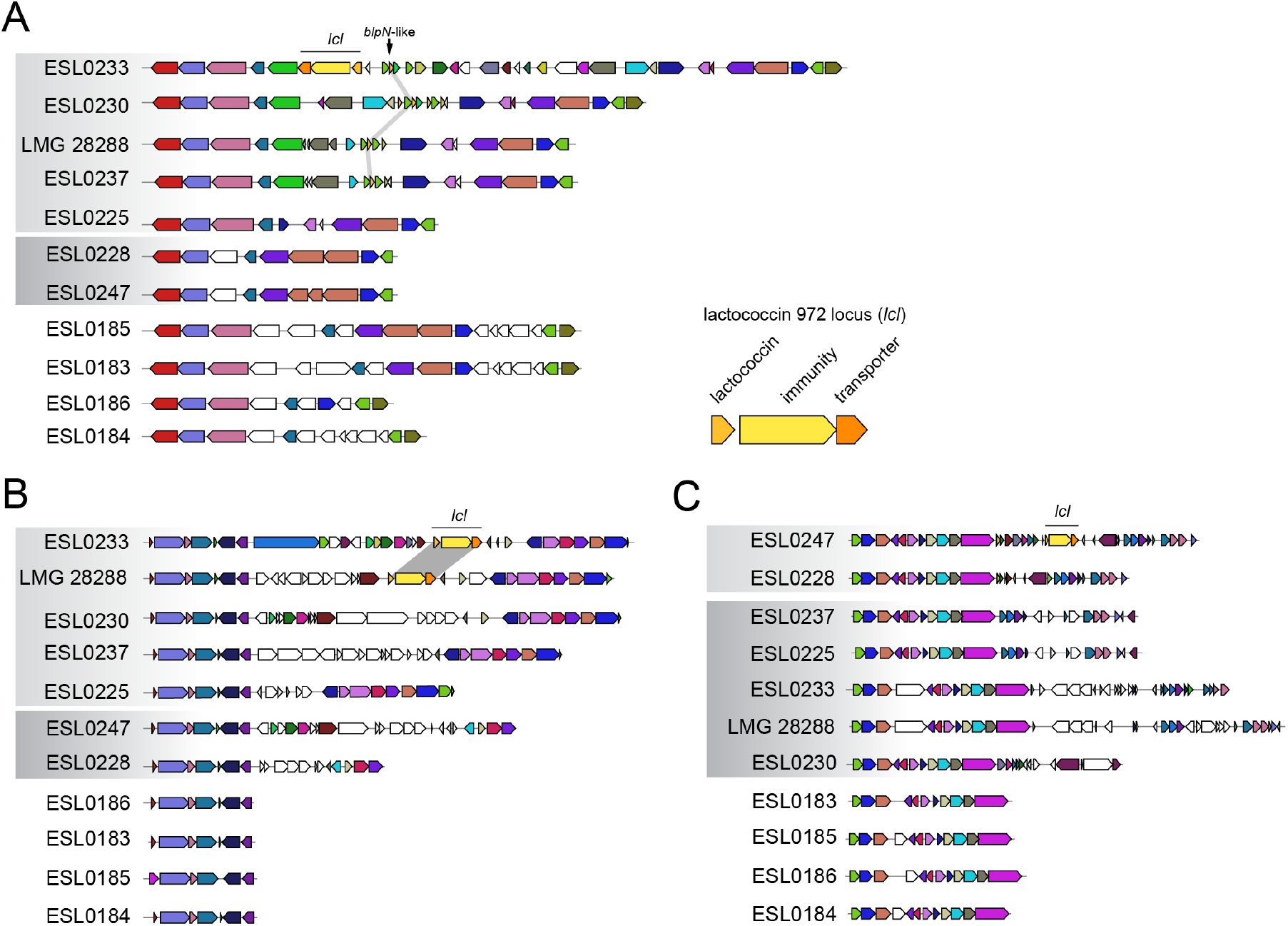
Additional genomic regions encoding class II bacteriocins in Firm5 strains of bumble bees. Genomic regions encoding bacteriocin genes were identified and visualized with MultiGeneBlast v1.1.14 (Medema *et al.* 2013). Arrows represent genes, and same color indicates homology. A black line indicates the lactococcin 972 locus (*lcl*) and vertical grey blocks connect the homologous genes in other strains. An enlarged version of the three genes of the *lcl* locus with annotation is shown in the lower right of panel A. Grey shading over strain names indicates two sublineages of bumble bee strains; the four honey bees strains are representatives of the four sublineages.

**Figure S9.**
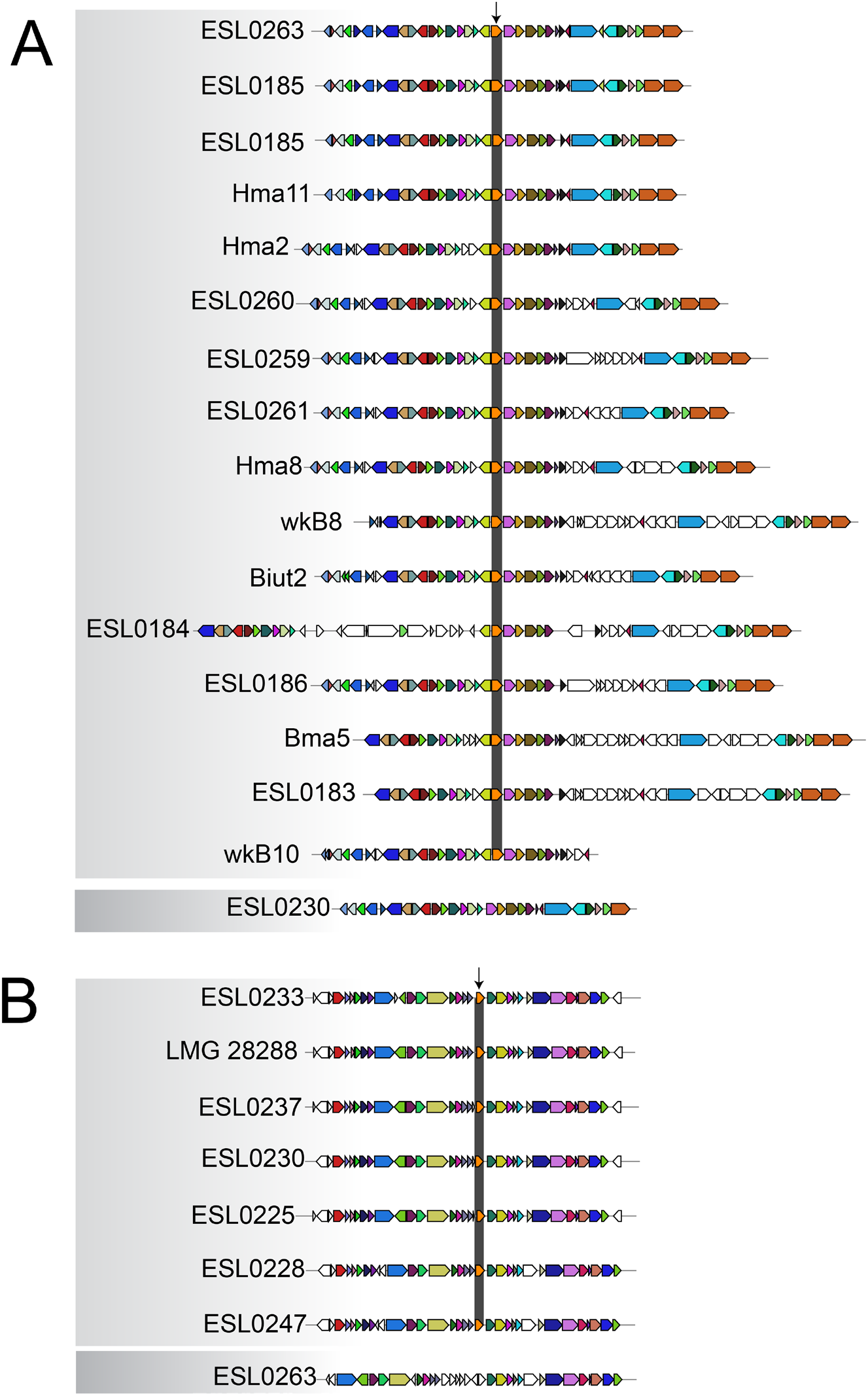
Genomic regions encoding helviticin-J, a class III bacteriocin. Genomic regions encoding bacteriocin genes were identified and visualized with MultiGeneBlast v1.1.14 (Medema *et al.* 2013). Arrows represent genes, and same color indicates homology. An arrow points at the helveticin-J gene homolog and vertical grey blocks connect the homologous genes in other strains. Strains with the two different types of grey shadings indicate strains from bumble bees and honey bees. **(A)** Genomic region encoding helveticin-J in honey bee strains, and **(B)** genomic region encoding helveticin-J in bumble bee strains.

## Supplementary Tables and Datasets (as separate files)

**Table S1.** Strain list and genome features.

**Table S2.** Pairwise 16S rRNA gene sequence identities.

**Table S3.** ANI values.

**Dataset S1.** List of gene families and their distribution according the three major groups: honey bee strains, bumble bee strains, outgroup strains.

**Dataset S2.** List of sublineage-specific gene families and COG category abbreviations.

**Dataset S3.** List of genes per genome with hits to the Carbohydrate-active enzyme (CAZY) database.

## References

Bankevich A, Nurk S, Antipov D et al. (2012) SPAdes: a new genome assembly algorithm and its applications to single-cell sequencing. Journal of computational biology: a journal of computational molecular cell biology, 19, 455–477.

Benjamini Y, Hochberg Y (1995) Controlling the false discovery rate: a practical and powerful approach to multiple testing. Journal of the Royal Statistical Society. Series B: Methodological 57:289–300.

Bobay L-M, Ochman H (2017) The Evolution of Bacterial Genome Architecture. Frontiers in genetics, 8, 829.

Bolger AM, Lohse M, Usadel B (2014) Trimmomatic: a flexible trimmer for Illumina sequence data. Bioinformatics, 30, 2114–20.

NCBI Resource Coordinators (2018) Database resources of the National Center for Biotechnology Information. Nucleic Acids Research, 46, D8–D13.

Corby-Harris V, Maes P, Anderson KE (2014) The bacterial communities associated with honey bee *(Apis mellifera)* foragers. PloS one, 9, e95056.

Cotter PD, Ross RP, Hill C (2013) Bacteriocins-a viable alternative to antibiotics? Nature reviews. Microbiology, 11, 95–105.

Cox-Foster DL, Conlan S, Holmes EC et al. (2007) A metagenomic survey of microbes in honey bee colony collapse disorder., 318, 283–287.

Duar RM, Frese SA, Lin XB et al. (2017) Experimental Evaluation of Host Adaptation of Lactobacillus reuteri to Different Vertebrate Species. Applied and environmental microbiology, 83, e00132–17.

Eddy SR (2009) A new generation of homology search tools based on probabilistic inference. Genome informatics. International Conference on Genome Informatics, 23, 205–211.

Ellegaard KM, Tamarit D, Javelind E et al. (2015) Extensive intra-phylotype diversity in lactobacilli and bifidobacteria from the honeybee gut. BMC genomics, 16, 284.

Ellegaard KM, Engel P (2019) Genomic diversity landscape of the honey bee gut microbiota. Nature communications, 10, 446.

Emery O, Schmidt K, Engel P (2017) Immune system stimulation by the gut symbiont *Frischella perrara* in the honey bee *(Apis mellifera)*. Molecular Ecology, 50, 735.

Engel P, Martinson VG, Moran NA (2012) Functional diversity within the simple gut microbiota of the honey bee. Proceedings of the National Academy of Sciences of the United States of America, 109, 11002–11007.

Eren AM, Sogin ML, Morrison HG et al. (2015) A single genus in the gut microbiome reflects host preference and specificity. The ISME journal, 9, 90–100.

Frese SA, Benson AK, Tannock GW et al. (2011) The Evolution of Host Specialization in the Vertebrate Gut Symbiont Lactobacillus reuteri (DS Guttman, Ed,). PLoS genetics, 7, e1001314.

Frese SA, MacKenzie DA, Peterson DA et al. (2013) Molecular Characterization of Host-Specific Biofilm Formation in a Vertebrate Gut Symbiont (DA Garsin, Ed,). PLoS genetics, 9, e1004057.

Guy L, Kultima JR, Andersson SGE (2010) genoPlotR: comparative gene and genome visualization in R. Bioinformatics, 26, 2334–2335.

Hebert PDN, Penton EH, Burns JM, Janzen DH, Hallwachs W (2004) Ten species in one: DNA barcoding reveals cryptic species in the neotropical skipper butterfly *Astraptes fulgerator*. Proceedings of the National Academy of Sciences, 101, 14812–14817.

Hutchinson GE (1957) Concluding remarks. Cold Spring Harbor Symposia on Quantitative Biology, 22: 415–427.

Jain C, Rodriguez-R LM, Phillippy AM, Konstantinidis KT, Aluru S (2018) High throughput ANI analysis of 90K prokaryotic genomes reveals clear species boundaries. Nature communications, 9, 5114.

Katoh K, Rozewicki J, Yamada KD (2017) MAFFT online service: multiple sequence alignment, interactive sequence choice and visualization. Briefings in bioinformatics, 30, 3059.

Kešnerová L, Mars RAT, Ellegaard KM et al. (2017) Disentangling metabolic functions of bacteria in the honey bee gut. PLoS biology, 15, e2003467.

Koch H, Abrol DP, Li J, Schmid-Hempel P (2013) Diversity and evolutionary patterns of bacterial gut associates of corbiculate bees. Molecular Ecology, 22, 2028–2044.

Kommineni S, Bretl DJ, Lam V et al. (2015) Bacteriocin production augments niche competition by enterococci in the mammalian gastrointestinal tract. Nature, 526, 719–722.

Kostic AD, Howitt MR, Garrett WS (2013) Exploring host-microbiota interactions in animal models and humans. Genes & development, 27, 701–718.

Kwong WK, Moran NA (2015) Evolution of host specialization in gut microbes: the bee gut as a model. Gut microbes, 6, 214–220.

Kwong WK, Moran NA (2016) Gut microbial communities of social bees. Nature reviews. Microbiology, 14, 374–384.

Kwong WK, Engel P, Koch H, Moran NA (2014) Genomics and host specialization of honey bee and bumble bee gut symbionts. Proceedings of the National Academy of Sciences of the United States of America, 111, 11509–11514.

Kwong WK, Medina LA, Koch H et al. (2017) Dynamic microbiome evolution in social bees. Science Advances, 3, e1600513.

Lee I, Kim YO, Park S-C, Chun J (2016) OrthoANI: An improved algorithm and software for calculating average nucleotide identity. International journal of systematic and evolutionary microbiology, 66, 1100–1103.

Letzel A-C, Pidot SJ, Hertweck C (2014) Genome mining for ribosomally synthesized and post-translationally modified peptides (RiPPs) in anaerobic bacteria. BMC genomics, 15, 983.

Ley RE, Hamady M, Lozupone C et al. (2008) Evolution of mammals and their gut microbes. Science, 320, 1647–1651.

Li L, Stoeckert CJ, Roos DS (2003) OrthoMCL: identification of ortholog groups for eukaryotic genomes. Genome research, 13,2178–89.

Ludvigsen J, Porcellato D, L’Abée-Lund TM, Amdam GV, Rudi K (2017) Geographically widespread honeybee-gut symbiont subgroups show locally distinct antibiotic-resistant patterns. Molecular Ecology, 26, 6590–6607.

Macarthur R, Levins R (2015) The Limiting Similarity, Convergence, and Divergence of Coexisting Species. The American Naturalist, 101, 377–385.

Markowitz VM, Chen I-MA, Chu K et al. (2014) IMG/M 4 version of the integrated metagenome comparative analysis system. Nucleic Acids Research, 42, D568–73.

Martinson VG, Moy J, Moran NA (2012) Establishment of characteristic gut bacteria during development of the honeybee worker. Applied and environmental microbiology, 78, 2830–2840.

Martinez B, Rodriguez A, Suárez JE (2000) Lactococcin 972, a bacteriocin that inhibits septum formation in lactococci. Microbiology, 146, 949–955.

McCutcheon JP, Moran NA (2012) Extreme genome reduction in symbiotic bacteria. Nature reviews. Microbiology, 10, 13–26.

McFall-Ngai M, Hadfield MG, Bosch TCG et al. (2013) Animals in a bacterial world, a new imperative for the life sciences. Proceedings of the National Academy of Sciences, 110, 3229–3236.

Medema MH, Takano E, Breitling R (2013) Detecting Sequence Homology at the Gene Cluster Level with MultiGeneBlast. Molecular Biolology Evolution, 30, 1218–23.

Moeller AH, Caro-Quintero A, Mjungu D et al. (2016) Cospeciation of gut microbiota with hominids. Science, 353, 380–382.

Moran NA, Hansen AK, Powell JE, Sabree ZL (2012) Distinctive gut microbiota of honey bees assessed using deep sampling from individual worker bees. PloS one, 7, e36393.

Ochman H, Worobey M, Kuo C-H et al. (2010) Evolutionary Relationships of Wild Hominids Recapitulated by Gut Microbial Communities. PLoS biology, 8, e1000546.

Oh PL, Benson AK, Peterson DA et al. (2010) Diversification of the gut symbiont Lactobacillus reuteri as a result of host-driven evolution. The ISME journal, 4, 377–387.

Olofsson TC, Alsterfjord M, Nilson B, Butler E, Vásquez A (2014) Lactobacillus apinorum sp. nov., Lactobacillus mellifer sp. nov., Lactobacillus mellis sp. nov., Lactobacillus melliventris sp. nov., Lactobacillus kimbladii sp. nov., Lactobacillus helsingborgensis sp. nov. and Lactobacillus kullabergensis sp. nov., isolated from the honey stomach of the honeybee Apis mellifera. International journal of systematic and evolutionary microbiology, 64, 3109–3119.

Praet J, Meeus I, Cnockaert M et al. (2015) Novel lactic acid bacteria isolated from the bumble bee gut: Convivina intestini gen. nov., sp. nov., Lactobacillus bombicola sp. nov., and Weissella bombi sp. nov. Antonie van Leeuwenhoek, 107, 1337–1349.

Rissman AI, Mau B, Biehl BS et al. (2009) Reordering contigs of draft genomes using the Mauve aligner. Bioinformatics, 25, 2071–2073.

Seedorf H, Griffin NW, Ridaura VK et al. (2014) Bacteria from diverse habitats colonize and compete in the mouse gut. Cell, 159, 253–266.

Sriswasdi S, Yang C-C, Iwasaki W (2017) Generalist species drive microbial dispersion and evolution. Nature communications, 8, 1162.

Stamatakis A (2014) RAxML version 8: a tool for phylogenetic analysis and postanalysis of large phylogenies. Bioinformatics, 30, 1312–1313.

Steele MI, Kwong WK, Whiteley M, Moran NA (2017) Diversification of Type VI Secretion System Toxins Reveals Ancient Antagonism among Bee Gut Microbes. mBio, 8, e01630–17.

Toft C, Andersson SGE (2010) Evolutionary microbial genomics: insights into bacterial host adaptation. Nature Reviews Genetics, 11, 465–475.

van Heel AJ, de Jong A, Montalban-Lopez M (2013). BAGEL3: automated identification of genes encoding bacteriocins and (non-)bactericidal posttranslationally modified peptides. Nucleic Acid Research, 41, W448–53.

Yin Y, Mao X, Yang J et al. (2012) dbCAN: a web resource for automated carbohydrate-active enzyme annotation. Nucleic Acid Research, 40, W445–51.

Zhang J, Kobert K, Flouri T, Stamatakis A (2014) PEAR: a fast and accurate Illumina Paired-End reAd mergeR. Bioinformatics, 30, 614–620.

Zheng H, Nishida A, Kwong WK et al. (2016) Metabolism of Toxic Sugars by Strains of the Bee Gut Symbiont Gilliamella apicola. mBio, 7, e01326–16.

Zheng J, Ruan L, Sun M, Gänzle M (2015) A Genomic View of Lactobacilli and Pediococci Demonstrates that Phylogeny Matches Ecology and Physiology. Applied and environmental microbiology, 81, 7233–7243.

